# Coupled protein quality control during nonsense mediated mRNA decay

**DOI:** 10.1101/2021.12.22.473893

**Authors:** Alison J. Inglis, Alina Guna, Ángel Gálvez Merchán, Akshaye Pal, Theodore K. Esantsi, Heather R. Keys, Evgeni M. Frenkel, Robert Oania, Jonathan S. Weissman, Rebecca M. Voorhees

## Abstract

Translation of mRNAs containing premature termination codons (PTCs) can result in truncated protein products with deleterious effects. Nonsense-mediated decay (NMD) is a surveillance pathway responsible for detecting and degrading PTC containing transcripts. While the molecular mechanisms governing mRNA degradation have been extensively studied, the fate of the nascent protein product remains largely uncharacterized. Here, we use a fluorescent reporter system in mammalian cells to reveal a selective degradation pathway specifically targeting the protein product of an NMD mRNA. We show that this process is post-translational, and dependent on an intact ubiquitin proteasome system. To systematically uncover factors involved in NMD-linked protein quality control, we conducted genome-wide flow cytometry-based screens. Our screens recovered known NMD factors, and suggested a lack of dependence on the canonical ribosome-quality control (RQC) pathway. Finally, one of the strongest hits in our screens was the E3 ubiquitin ligase CNOT4, a member of the CCR4-NOT complex, which is involved in initiating mRNA degradation. We show that CNOT4 is involved in NMD coupled protein degradation, and its role depends on a functional RING ubiquitin ligase domain. Our results demonstrate the existence of a targeted pathway for nascent protein degradation from PTC containing mRNAs, and provide a framework for identifying and characterizing factors involved in this process.

## INTRODUCTION

Nonsense mediated mRNA decay (NMD) is a broadly conserved and essential surveillance pathway that ensures the integrity of the transcriptome and regulates the levels of many cellular mRNA transcripts. NMD was initially identified for its role in recognizing and degrading aberrant, disease-causing mRNAs that contain a premature termination codon (PTC) within their open reading frame (Chang & Kan, 1979; Losson & Lacroute, 1979; Maquat et al., 1981). When translated, these mRNAs produce truncated proteins that can be aggregation-prone, develop gain of function phenotypes, (Nonaka et al. 2009) or have dominant negative effects (Dietz et al., 1993; Hall & Thein, 1994; Kugler et al., 1995; Thein et al., 1990). NMD thus plays a critical role in maintaining cellular proteostasis by preventing expression of these potentially deleterious truncated proteins. Further, one third of genetic disorders (Mort et al., 2008), including muscular dystrophy (Kerr et al., 2001) and cystic fibrosis (O’Sullivan, 2014) and many cancers (Anczuków et al., 2008; Karam et al., 2008; Perrin-Vidoz et al., 2002; Reddy et al., 1995; Ware et al., 2006) are the result of PTC-causing mutations that lead to recognition and degradation of the resulting mRNAs by NMD.

In addition to its role in transcriptome maintenance, NMD also regulates the levels of roughly 10% of endogenous transcripts, facilitating rapid and flexible changes in gene expression in response to environmental and developmental stimuli (He et al., 2003; Lelivelt & Culbertson, 1999; Rehwinkel et al., 2005). NMD thus plays a fundamental role in diverse, but physiologically essential processes including regulating the temporal expression of proteins during the cell cycle (Choe et al., 2014); degrading PTC-containing transcripts produced by somatic recombination during immune system development (Bruce & Wilkinson, 2003); and suppressing viral gene expression as a component of the innate immune response (Balistreri et al., 2014; Ramage et al., 2015).

While there are no definitive rules as to what defines an NMD substrate, the composition of protein factors that decorate the 3’UTR of an mRNA seem to either promote or prevent its degradation via NMD (Behm-Ansmant et al., 2007; Singh et al., 2008). For example, the positioning of poly(A) binding protein (PABP) adjacent to the termination codon has been shown to be protective (Silva et al., 2008), while unusual physical features such as upstream open reading frames (uORFs) and long 3’UTRs are established cues for degradation by NMD (Behm-Ansmant et al., 2007; Mendell et al., 2004; Singh et al., 2008). It has also been observed that the many apparently ‘normal’ transcripts that are regulated by NMD have lower codon optimality and a higher rate of out-of-frame translation (Celik et al., 2017). However, the best characterized trigger for recognition by NMD is the presence of an intron downstream of a stop codon, which is commonly the result of genetic mutations or defects in alternative splicing (Shoemaker & Green, 2012). Splicing of these introns results in the deposition of an exon-junction complex (EJC) 24 nucleotides upstream of the splice site, which is retained upon packaging and export to the cytoplasm (Ballut et al., 2005; Hoskins & Moore, 2012; le Hir et al., 2000, 2001). Because the majority of endogenous stop codons are localized within the last exon of protein coding genes, EJCs are typically removed during translational elongation (Dostie & Dreyfuss, 2002). The persistence of an EJC downstream of a stop codon is thus a characteristic of a PTC-containing mRNA and results in robust recognition by the NMD pathway (Gehring et al., 2003; Palacios et al., 2004).

Translation termination in the presence of a downstream EJC triggers NMD through a network of interactions between the core NMD factors UPF1, UPF2 and UPF3B; the downstream EJC; and the translational termination factors including eRF1 and eRF3 (Chamieh et al., 2008; Czaplinski et al., 1998; Kim et al., 2001; le Hir et al., 2001). Phosphorylation of UPF1 by SMG1 recruits a suite of RNA decay machinery to decap (DCP2) (Cho et al., 2009; Lai et al., 2012), deadenylate (CCR4-NOT) (Loh et al., 2013), cleave (SMG6) (Eberle et al. 2009; Huntzinger et al. 2008), and ultimately degrade the associated mRNA.

Like other mRNA surveillance pathways, NMD substrates are recognized and targeted for degradation co-translationally (Belgrader et al., 1993; J. Wang et al., 2002; Zhang & Maquat, 1997), resulting in the synthesis of a potentially aberrant nascent polypeptide chain. Pathways such as no-go and non-stop mRNA decay rely on a coordinated protein quality control pathway, known as ribosome associated quality control (RQC) to both rescue the ribosome and concomitantly target the nascent protein for degradation (Doma & Parker, 2006; Frischmeyer et al., 2002; Juszkiewicz et al., 2018; van Hoof et al., 2002). In both cases, a terminally stalled ribosome or a collided diribosome triggers ribosome splitting (Becker et al., 2011; Pisareva et al., 2011; Shao et al., 2015, 2016; Shoemaker & Green, 2012) and nascent chain ubiquitination by the E3 ligase LTN1 (facilitated by NEMF, TAE2, and P97) (Brandman et al., 2012; Defenouillère et al., 2013; Lyumkis et al., 2014; Shao et al., 2013, 2015; Verma et al., 2013). The ubiquitinated nascent chain is then released from the ribosome by the endonuclease ANKZF1 (Vms1 in yeast) for degradation by the proteasome (Rendón et al., 2018; Verma et al., 2018).

Given the potential dominant negative and proteotoxic effects of even small amounts of a truncated NMD substrate, it has been suggested that a similar protein quality control pathway may exist to recognize and degrade nascent proteins that result from translation of NMD mRNAs. Indeed, proteins produced from PTC-containing mRNAs are less stable than those from normal transcripts (Kuroha, Tatematsu, and Inada 2009; Kuroha et al. 2013; Pradhan et al. 2021; Udy and Bradley 2021). However, these observations are largely based on comparison of truncated products with longer, potentially more stable polypeptides, making it difficult to distinguish NMD linked protein degradation from general cellular quality control mechanisms. Recent studies more directly test this using a full-length protein product, but have not defined the mechanism of its targeting and degradation, nor directly identified a role for the ubiquitin-proteasome pathway. Furthermore, though it has been postulated that components of the RQC are involved in turnover of nascent NMD substrates (Arribere & Fire, 2018), the factors required for this process have not been systematically investigated. Because NMD is triggered at a stop codon unlike no-go and non-stop decay, a putative NMD-coupled protein quality control pathway may require a fundamentally different strategy to initiate nascent protein degradation.

Here we describe a reporter system that we have used to definitively define and characterize a coupled protein quality control branch of NMD. We demonstrate that in addition to triggering mRNA degradation, NMD concomitantly coordinates degradation of the nascent polypeptide via the ubiquitin-proteasome pathway. Using this reporter system, we systematically identify factors required for NMD-coupled protein degradation, which are distinct from the canonical rescue factors of the RQC. Characterization of a coupled protein-degradation branch of NMD represents a new facet of our understanding of how the cell ensures the integrity and composition of its proteome, and sheds further light on the interplay between mRNA and protein quality control.

## RESULTS

### A reporter strategy to decouple mRNA and protein quality control in NMD

To identify a putative NMD-linked protein quality control pathway, we developed a reporter system that sought to uncouple mRNA and protein quality control during NMD. The reporter consists of a single open reading frame expressing GFP and RFP, separated by a viral 2A sequence that causes peptide skipping (Y. Wang et al., 2015) (Fig. 1A, sFig. 1A). A robust example of an endogenous NMD substrate is the β-globin gene with a nonsense mutation at codon 39, which results in a premature stop codon followed by an intron (Jing et al., 1998). We therefore reasoned that positioning the first intron of the human β-globin gene into the 3’UTR of our reporter after the stop codon would also lead to its recognition as an NMD substrate, as has been previously reported (Chu et al., 2021; Lykke-Andersen et al., 2000; Pereverzev et al., 2015). We confirmed that the exogenous β-globin intron is efficiently spliced (sFig.1B), and observed that the mRNA levels of the NMD reporter were ∼5-fold lower than a matched non-NMD control (Fig. 1B). We found that the GFP fluorescence of the NMD reporter and control correlated with their respective mRNA levels, as directly measured by qPCR, suggesting that GFP fluorescence can be used as a proxy for transcript levels (sFig. 1C). Finally, knockdown of the core NMD factor UPF1 specifically increased the GFP fluorescence of the NMD reporter (sFig. 1F), but had no effect on the matched control. We therefore concluded that our fluorescent reporter is recognized and degraded in an NMD-dependent manner.

**Figure 1.**
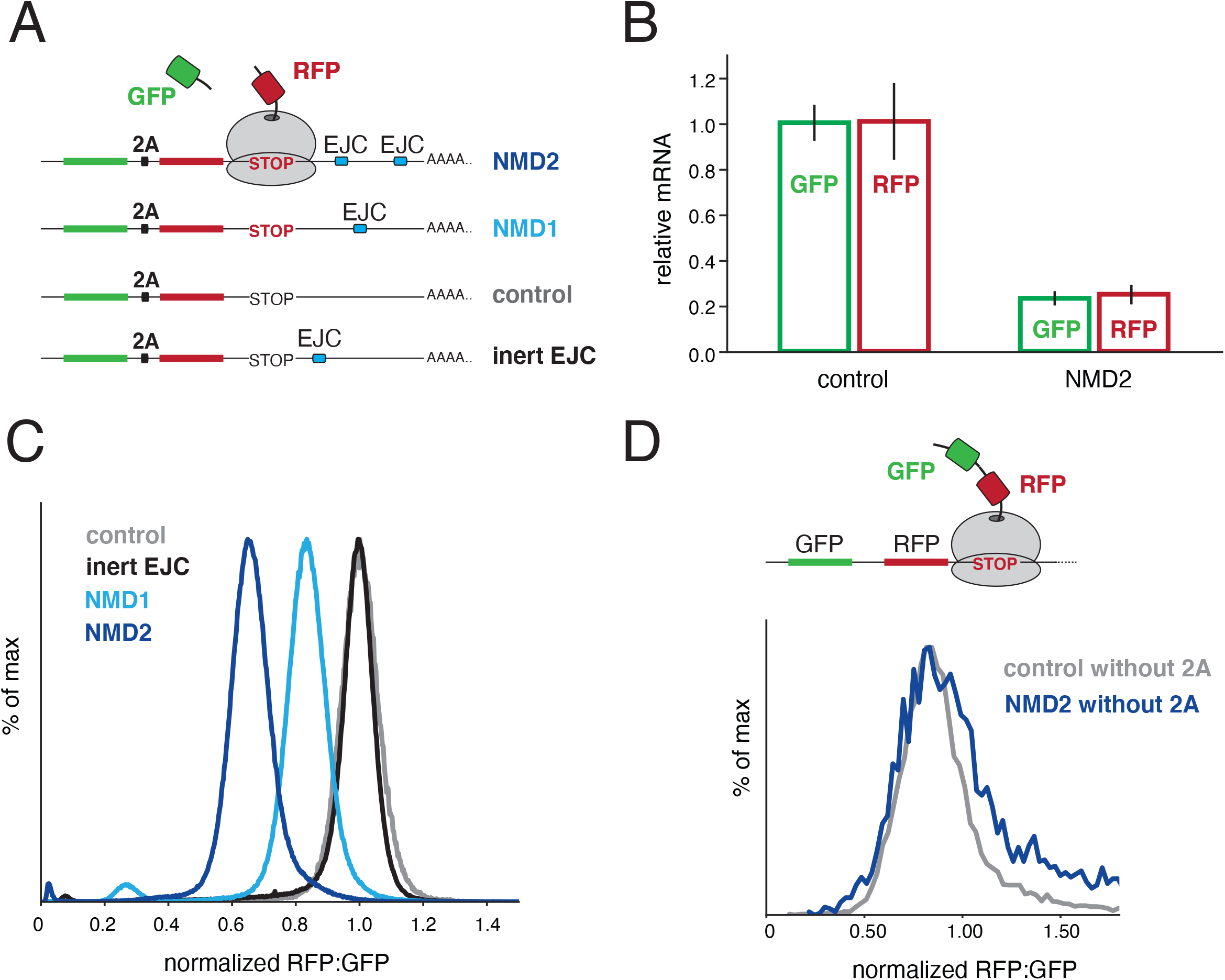
Destabilization of nascent proteins from PTC-containing mRNAs. (A) Schematic of the reporter strategy used to decouple protein and mRNA degradation in NMD. GFP and RFP are encoded in a single open reading frame separated by a viral 2A sequence. Positioning an intron within the 3’ UTR results in deposition of an exon junction complex (EJC) upon splicing, trigger-ing NMD when compared to a matched control (stop codon depicted in red). Either one or two in-trons derived from the β-globin gene are inserted after the stop codon (NMDl and NMD2 respectively). To control for the documented stimulation in translation that results from the presence of an EJC (Nott et al., 2004), we created a reporter in which the intron was positioned twelve nucleotides after the stop codon, a distance insufficient for recognition as an NMD substrate (inert EJC) (Nagy & Maquat, l998). (B) Stable cell lines expressing either the control or the reporter was induced with doxycycline for 24 hours and the total mRNA was then purified. Relative mRNA levels were determined by RT-qPCR using primers that anneal to the very 5’ region of the GFP and 3’ region of the RFP open reading frames. The results were normalized to the control and the standard deviation from three independent experiments is displayed. (C) Stable cell lines for the indicated report-ers were analyzed by flow cytometry. The ratio of RFP:GFP fluorescence, as normalized to the control reporter is depicted as a histogram. (D) The NMD2 and control reporters in which the 2A sequence was scrambled, resulting in tethering of both GFP and RFP to the ribosome at the stop codon were analyzed as in (C).

Recognition of our reporter as an NMD substrate, and subsequent mRNA decay, is a pre-requisite for establishing whether there is an additional pathway dedicated to nascent protein degradation. To address this, our reporter has two important physical features. First, it can be used to deconvolute post-transcriptional versus post-translational effects on reporter fluorescence. Upon translation, the GFP is released by the 2A sequence while the RFP remains tethered to the ribosome until the termination codon, where NMD is initiated by interaction between the downstream EJC and the ribosome. We reasoned that if there is an NMD-coupled pathway that triggers degradation of the nascent polypeptide, it would thus act only on the RFP but not the released GFP, resulting in a reduction in the RFP:GFP ratio in comparison to a matched control. In contrast, if NMD functions only in mRNA degradation, we would expect a decrease in both the RFP and GFP levels but would observe no change in the RFP:GFP ratio. Second, these reporters can specifically distinguish nascent protein degradation by a coupled protein quality control pathway from non-specific recognition by general cellular quality control machinery. Canonical NMD substrates contain PTCs that result in translation of a truncated protein, which may be misfolded and thus recognized and degraded by non-specific cytosolic quality control pathways (Popp & Maquat, 2013). By instead using an intact RFP moiety that is recognized as an NMD substrate only because of an intron in its 3’UTR, any destabilization of RFP must result from a coordinated event that occurs prior to its release from the ribosome.

Indeed, using flow cytometry, we observed a decrease in RFP:GFP fluorescence for an NMD substrate compared to a matched control (Fig. 1C). Addition of a second β-globin intron to the 3’UTR (Hoek et al., 2019) resulted in a larger decrease in both the mRNA levels and RFP:GFP fluorescence, suggesting the two effects may be tightly coordinated (Hoek et al. 2019). While this decrease in RFP:GFP levels was consistent with NMD-dependent protein quality control, we sought to exclude several alternative models that could also account for this observation. First, we swapped the order of the RFP and GFP to rule out that differential maturation and/or turnover rates of the fluorophores could explain the decrease in RFP:GFP ratio (sFig. 1D) (Amrani et al., 2004; Balleza et al., 2018). Second, we considered whether the decrease in RFP:GFP ratio could be the result of NMD-dependent deadenylation and 3’ to 5’ exonuclease degradation of the reporter mRNA (Chen & Shyu, 2003; Mitchell & Tollervey, 2003; Takahashi et al., 2003). However, we detected no change in the relative mRNA levels of the RFP and GFP coding regions of the NMD substrate (Fig. 1B), confirming that the effect must occur post-transcriptionally.

Finally, we addressed two related possibilities: whether slow translational termination, characteristic of NMD substrates (Amrani et al. 2004), or SMG6-dependent endonucleolytic cleavage of the mRNA at the stop codon could explain the RFP:GFP ratio decrease (Eberle et al. 2009). The former would result in increased dwell time of the ribosome at the stop codon when the ∼30 C-terminal residues of RFP remain occluded in the ribosomal exit tunnel and could prevent RFP fluorescence. The latter would lead to production of full-length GFP but truncated RFP, and would be consistent with models proposed for putative NMD-coupled protein quality control in *C. elegans* (Arribere & Fire, 2018). However, appending a flexible linker to the C-terminus of RFP to ensure it is fully emerged from the ribosome at the stop codon did not affect the RFP:GFP ratio (Fig. 1D). Conversely, scrambling the 2A sequence, such that both the GFP and RFP are tethered to the ribosome at the stop codon, abolished the ratio difference (Fig. 1E). Together these data exclude that the NMD-dependent decrease in RFP:GFP ratio is due to changes in translation rate, processivity, peptide release, endonucleolytic cleavage, or preferential 3’-5’ degradation.

### NMD-dependent protein degradation occurs via the ubiquitin proteasome pathway

Having established that an NMD-dependent decrease in RFP fluorescence occurs post-translationally, we tested whether inhibition of the ubiquitin-proteasome pathway could rescue the observed phenotype. We found that both the proteasome inhibitor MG132 and the E1 ubiquitin-activating enzyme inhibitor MLN7243 specifically increased the RFP:GFP ratio of the NMD reporter (Fig. 2A; sFig. 2A). Importantly, this increase was due to an effect on RFP and not GFP (Fig. 2B), consistent with the model that NMD-dependent protein degradation acts post-translationally and selectively toward the polypeptide associated with the ribosome at the PTC.

**Figure 2.**
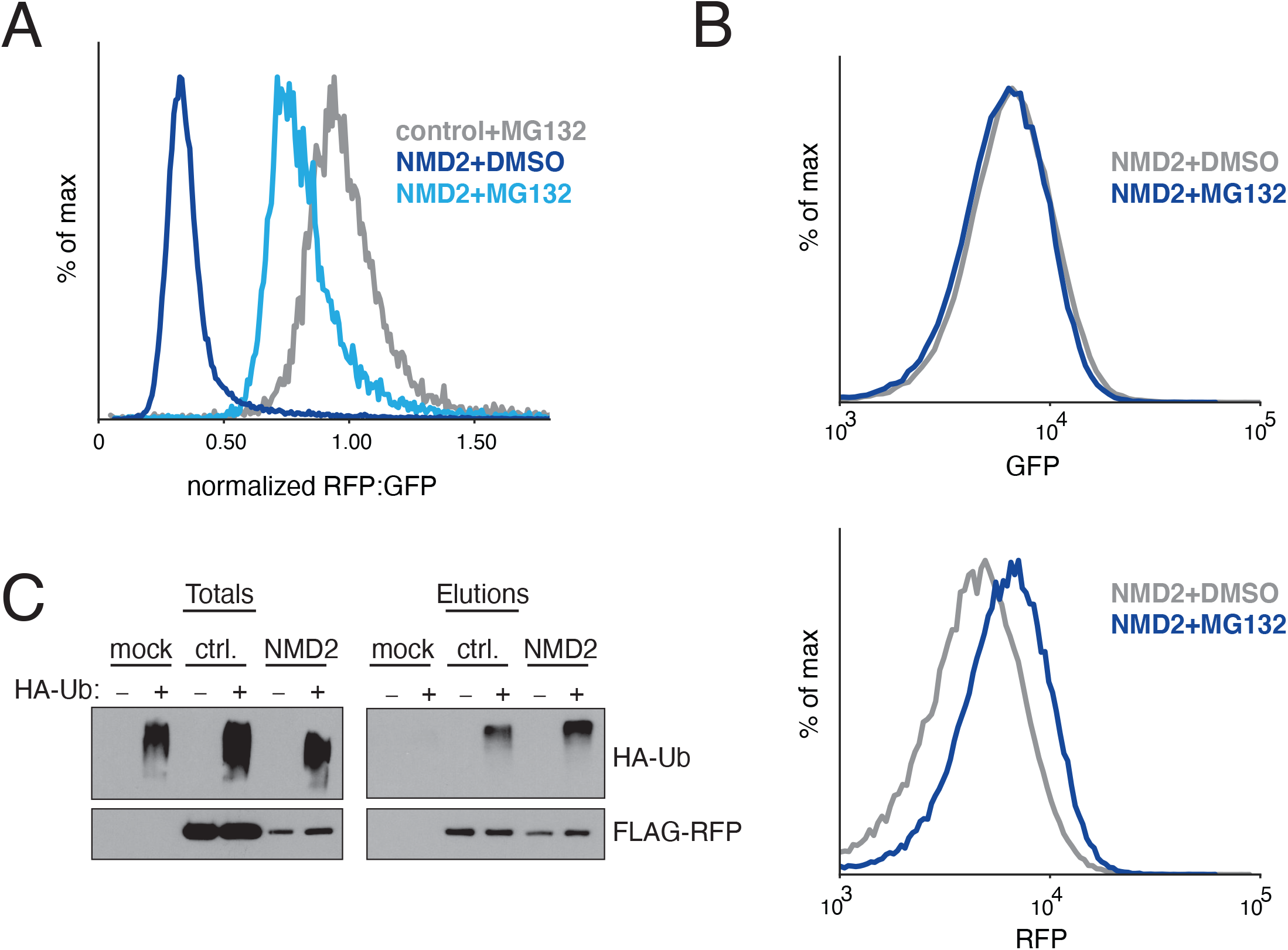
NMD-dependent protein degradation occurs via the ubiquitin proteasome pathway. (A) Flow cytometry analysis of HEK293T cells transiently transfected with either the control or NMD2 reporter (Fig. 1A) and treated with the proteasome inhibitor MG132 or DMSO. (B) As in (A) using stable K562 cell lines expressing an inducible NMD2 reporter treated with either MG132 or DMSO. Shown are the GFP (top) and RFP (bottom) channels for the indicated condi-tions displayed as a histogram. (C) HEK-293T cells were transiently transfected with either the control or NMD2 reporter (modified to incorporate a 3xFLAG tag at the N-terminus of RFP) in the presence of HA-tagged ubiquitin (HA-Ub). To stabilize ubiquitinated species, cells were treated with MG132 prior to lysis, and RFP was immunoprecipitated with anti-FLAG resin. Ubiquitinated species were detected by Western blotting for HA-Ub.

To confirm that the observed changes in fluorescence reflect changes at the protein level, we directly tested for stabilization of RFP upon E1 enzyme inhibition by Western blotting (sFig. 2B). The absence of truncated RFP would be consistent with a model in which NMD-dependent protein quality control is initiated at the stop codon. Finally, we directly observed a marked increase in ubiquitination of nascent NMD substrates compared to a matched control, excluding potential indirect effects of ubiquitin-proteasome pathway inhibition (Fig. 2C). Therefore, we concluded that in addition to its well-characterized role in mRNA degradation, NMD also triggers degradation of nascent proteins via the ubiquitin proteasome pathway.

### Identification of factors required for NMD-coupled protein quality control

Using our characterized reporter, we systematically identified factors required for the protein degradation arm of NMD using a fluorescence-activated cell sorting (FACS) based CRISPR interference (CRISPRi) (Horlbeck et al., 2016) and CRISPR knockout (CRISPR-KO) screen. We reasoned that the knockdown screen would enable study of essential proteins, including the core NMD factors UPF1 and UPF2 (Hart et al., 2017). Conversely, the knockout screen would identify factors that require near-complete depletion to induce a measurable phenotype, which can lead to false negatives in CRISPRi screens (Rosenbluh et al., 2017). To do this, we engineered two K562 human cell lines that expressed the NMD2 reporter either alone or with the CRISPRi silencing machinery (Gilbert et al., 2014). We transduced the CRISPRi cell line with a single guide RNA (sgRNA) library targeting all known protein-coding open reading frames as previously described (hCRISPRi-v2) (Horlbeck et al., 2016). For the knockout screen, we used a novel 100,000 element library that targets all protein encoding genes (∼5 sgRNA/gene), which we used to simultaneously deliver both the genome wide sgRNA library and cas9 (see methods).

We hypothesized that depletion of factors required for NMD-coupled protein quality control would stabilize RFP, thereby increasing the RFP:GFP ratio. However, depletion of factors that impede NMD-coupled protein quality control would further decrease the RFP:GFP ratio. For the CRISPRi screen, after eight days of knockdown, we sorted cells with high and low RFP:GFP ratios via FACS, and identified sgRNAs enriched in those cells by deep sequencing. For the knockout screen we isolated cells with perturbed RFP:GFP ratios on days eight, ten and twelve post infection of the CRISPR-KO library. We postulated that essential genes would be better represented at the earlier time points before their depletion becomes lethal, while factors that require complete depletion and/or have longer half-lives would be detected at later time points.

In both the knockdown and knockout screens, we find substantial differences between the hits identified here and those reported from earlier NMD RNA-degradation screens (Alexandrov et al., 2017; Baird et al., 2018; Sun et al., 2011; Zinshteyn et al., 2021), suggesting our reporter is indeed specific to the protein quality control branch of NMD. However, we also identified several splicing and core NMD factors as effectors of the RFP:GFP ratio. For example, we found that the core component of the EJC, CASC3 (Gerbracht et al., 2020) is required for NMD-coupled protein degradation (Fig. 3B, 3C). Furthermore, depletion of several known NMD factors—UPF1, UPF2, UPF3b, SMG6—increased the RFP:GFP ratio of our NMD-reporter. Together, these results suggest a single, shared recognition step for both the mRNA and protein quality control branches of NMD, which requires recognition of an intact EJC downstream of the stop codon via interactions between the canonical NMD factors and the ribosome.

**Figure 3.**
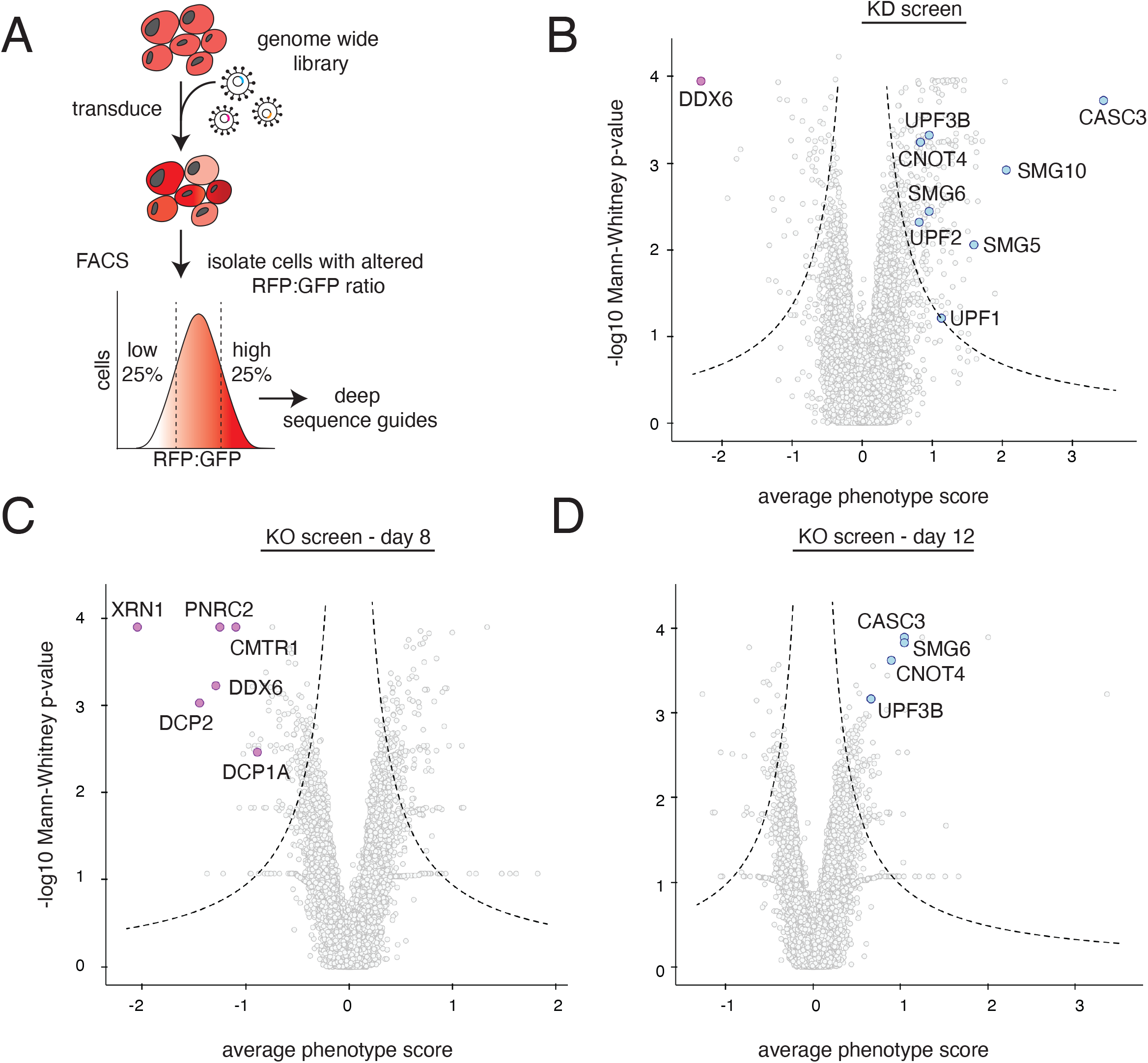
Systematic characterization of factors required for NMD-coupled protein quality control. (A) Schematic of the workflow used to carry-out the FACS-based reporter screens to identify factors involved in NMD-linked nascent chain degradation. K562 cells reporter cell lines contained a tet-inducible NMD2 reporter and were infected with either a whole-genome CRISPRi sgRNA library or a CRISPR-KO library. Reporter expression was induced with doxycycline for 24 hours prior to cell sorting. Cells were sorted based on ratiometric changes in RFP relative to GFP, and the sgRNA expressed in those cells were identified using deep sequencing. The CRISPR knockout screen was sorted on days 8, 10 and 12 post library infection to account for drop out of essential genes. The CRISPRi screen was sorted on day 8. (B) Volcano plot of the RFP:GFP stabilization phenotype (log2 for the three strongest sgRNAs) and Mann-Whitney p values from the genome-wide CRISPRi screen. Genes falling outside the dashed lines are statisti-cally significant. Each gray point represents a gene. Notable hits causing an increase in the RFP to GFP ratio are shown in light blue and include known NMD factors (UPF1, UPF2, UPF3B, SMG5, SMG6. SMG10) and the E3 ligase CNOT4. DDX6, a known suppressor of NMD, which causes a lower RFP to GFP ratio, is shown in purple. (C) Volcano plot as in (B) from the ge-nome-wide CRISPR knock-out screen sorted at an early time point, prior to essential gene drop out. In purple are highlighted factors that cause a decrease in RFP relative to GFP. These include genes involved in mRNA decapping (PNRC1, CMTR1, DCP1A, and DCP2), DDX6, and the 5’-3’ exonuclease XRN1. (D) As in (C) but from day 12. In blue are shown known NMD factors (CASC3, SMG6, UPF3B) and the E3 ligase CNOT4.

At day eight of the knockout screen, we found that several essential factors required for 5’ to 3’ mRNA degradation were enriched in the population of cells with lower RFP:GFP fluorescence (Fig. 3C). In both the knockdown and knockout screen, we found that depletion of the E3 ubiquitin ligase CNOT4 increased the RFP:GFP ratio of the reporter, suggesting a potential role in NMD-coupled protein quality control.

### NMD-coupled protein quality control is not mediated by canonical RQC factors

Notably absent in both the knockdown and knockout screen were canonical components of the RQC pathway, suggesting that NMD substrates may rely on an alternative strategy for nascent protein degradation. Because the CRISPRi screen was performed using the same strategy and conditions as earlier reporter screens for non-stop decay—including the same cell type, sgRNA library, and sampling time point—the screens are directly comparable (Hickey et al., 2020). While depletion of RQC factors including pelota and the E3 ubiquitin ligase LTN1 were identified in the non-stop reporter screen, neither are significant hits for NMD-dependent protein degradation in our system (Fig. 4A, 4B). We directly verified that LTN1 knockdown has no effect on our NMD reporter, though had a marked effect on the fluorescence ratio of an established non-stop decay substrate (Fig. 4C). We therefore concluded that NMD-coupled protein degradation is mediated by a new, uncharacterized set of factors (Chu et al., 2021).

**Figure 4.**
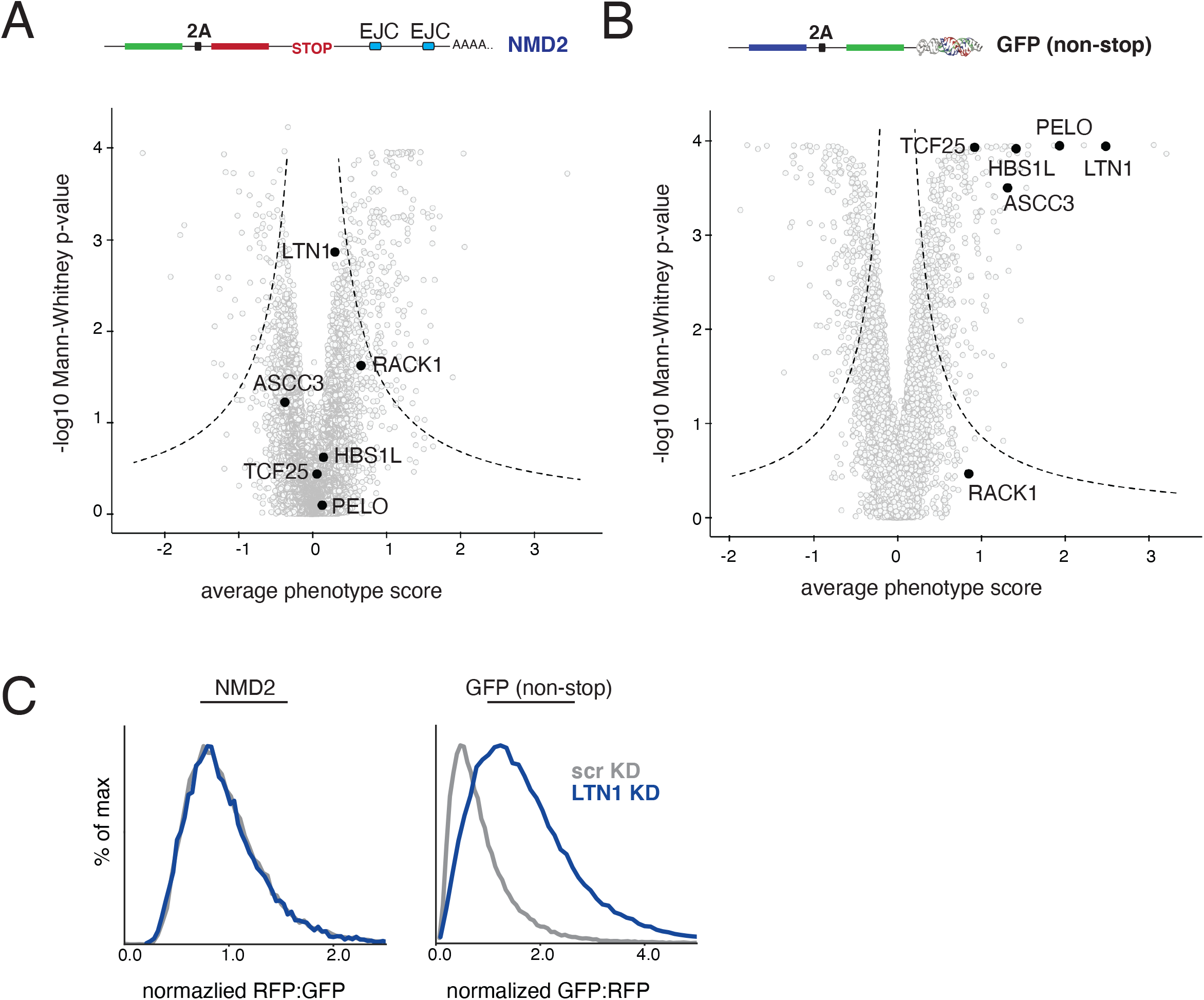
NMD-linked protein degradation is not mediated by the canonical RQC pathway. (A) Volcano plot of the NMD2 reporter CRISPRi screen as in Fig 3A. Highlighted in black are factors involved in the canonical RQC. (B) For comparison, RQC factors are highlighted in black on a volcano plot for a earlier CRISPRi screen using a non-stop reporter conducted using identical conditions as in (A) (Hickey et al., 2020). (C) K562 cells containing CRISPRi machinery and either an inducible NMD2 reporter or a constitutively expressed bidirectional GFP non-stop reporter (as in Hickey, 2020) were infected with a sgRNA targeting the E3 ligase LTN1. The RFP to GFP ratio for NMD2, and the GFP to RFP ratio for the GFP non-stop reporter as determined by flow cytometry are displaced as a histogram.

### Factors required for NMD-coupled protein quality control

Hits from the FACS based reporter screens were validated using an arrayed screen with a matched non-NMD control. These data confirmed that knockdown of the splicing factor CASC3 increased both the GFP levels and the RFP:GFP ratio of our NMD reporter (Fig. 5A). Knockdown of the 5’decapping enzyme DCP1A also increased GFP levels, but decreased the RFP:GFP ratio.

**Figure 5.**
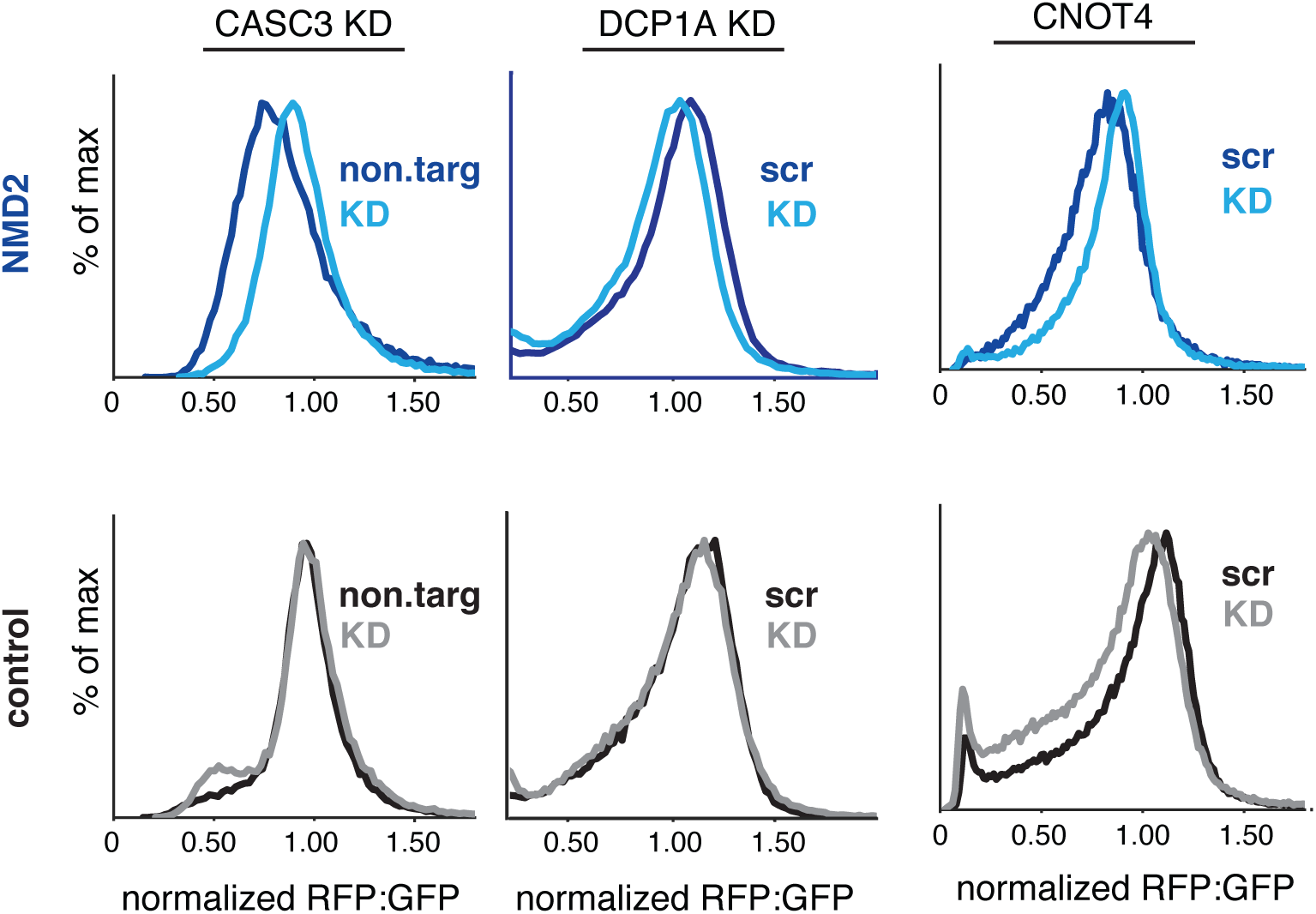
Validation of factors involved in NMD-coupled protein quality control. Panel of validated hits from NMD2 screens. Factors of interest were depleted by either sgRNA in K562 cells, or by siRNA in HEK293T cells as indicated. Displayed are the RFP:GFP ratio for the NMD2 (top) and control (bottom) reporters as determined by flow cytometry. Data was normalized to the scrambled siRNA (scr) or the non-targeting guide controls (non.targ).

Having observed that the nascent protein is directly ubiquitinated and degraded by the proteasome (Fig 2), we were particularly interested in identifying an E3 ubiquitin ligase responsible for targeting the NMD-linked nascent chain for degradation. The core NMD factor UPF1 is an E3 ubiquitin RING ligase (Takahashi et al., 2008) and thus would be well-positioned to mediate nascent chain degradation during NMD. Previous studies have demonstrated that UPF1 stimulates proteasomal degradation of proteins expressed from NMD-targeted mRNA transcripts in yeast, with reporter stability significantly increased in *upf1* knockout strains; however, the mechanism underlying this phenotype is unclear and a direct role in nascent chain ubiquitination by UPF1 was not shown (Kuroha et al., 2009). UPF1 was identified as a weak hit in our CRISPRi screen (Fig. 3B), and its depletion resulted in a shift in the RFP:GFP ratio of the NMD reporter (sFig. 1F). However, rescue of UPF1 knockdown with a RING mutant that disrupts binding with E2 ubiquitin-conjugating enzymes (Feng et al., 2017) phenocopied wild-type UPF1 in restoring both the GFP levels and RFP:GFP ratio of our NMD reporter (sFig. 3). This result would be inconsistent with a role for the RING domain of UPF1 in ubiquitination of the nascent protein, and suggests that the involvement of UPF1 may instead be upstream of the protein degradation branch.

In addition to UPF1, we identified four additional E3 ubiquitin ligases in either the knock-down and knockout screen (KEAP1, MYLIP, CBLL1, and TRIM25). However, the only ligase identified in both screens was the RING ligase CNOT4 (Fig 3B, 3C). CNOT4 is a conserved component of the multi-subunit CCR4-NOT complex, which regulates eukaryotic gene expression by deadenylation, i.e. processive shortening of mRNA poly(A) tails (Collart, 2016; Yamashita et al., 2005). CNOT4 is not a core structural component of the CCR4-NOT complex, is not required for deadenylation, and in mammals a population of CNOT4 exists independently from the rest of the CCR4-NOT complex (Jeske et al., 2006; Lau et al., 2009). Indeed, other members of the CCR4-NOT complex were not identified as significant hits in either the knockdown or knockout NMD screens (sFig. 4A). However, the matched CRISPRi screen for non-stop decay identified CNOT1 as a robust hit, verifying the efficacy of the sgRNAs (Hickey et al, 2021; sFig. 4B). Further, CNOT4 contains an N-terminal RING ligase domain (Albert et al., 2002; Hanzawa et al., 2001), as well as a conserved RNA-binding motif and zinc finger domain (Fig. 6A) (Inada and Makino 2014; Panasenko 2014). These data together suggest an independent role for CNOT4 in NMD-linked nascent protein degradation.

**Figure 6.**
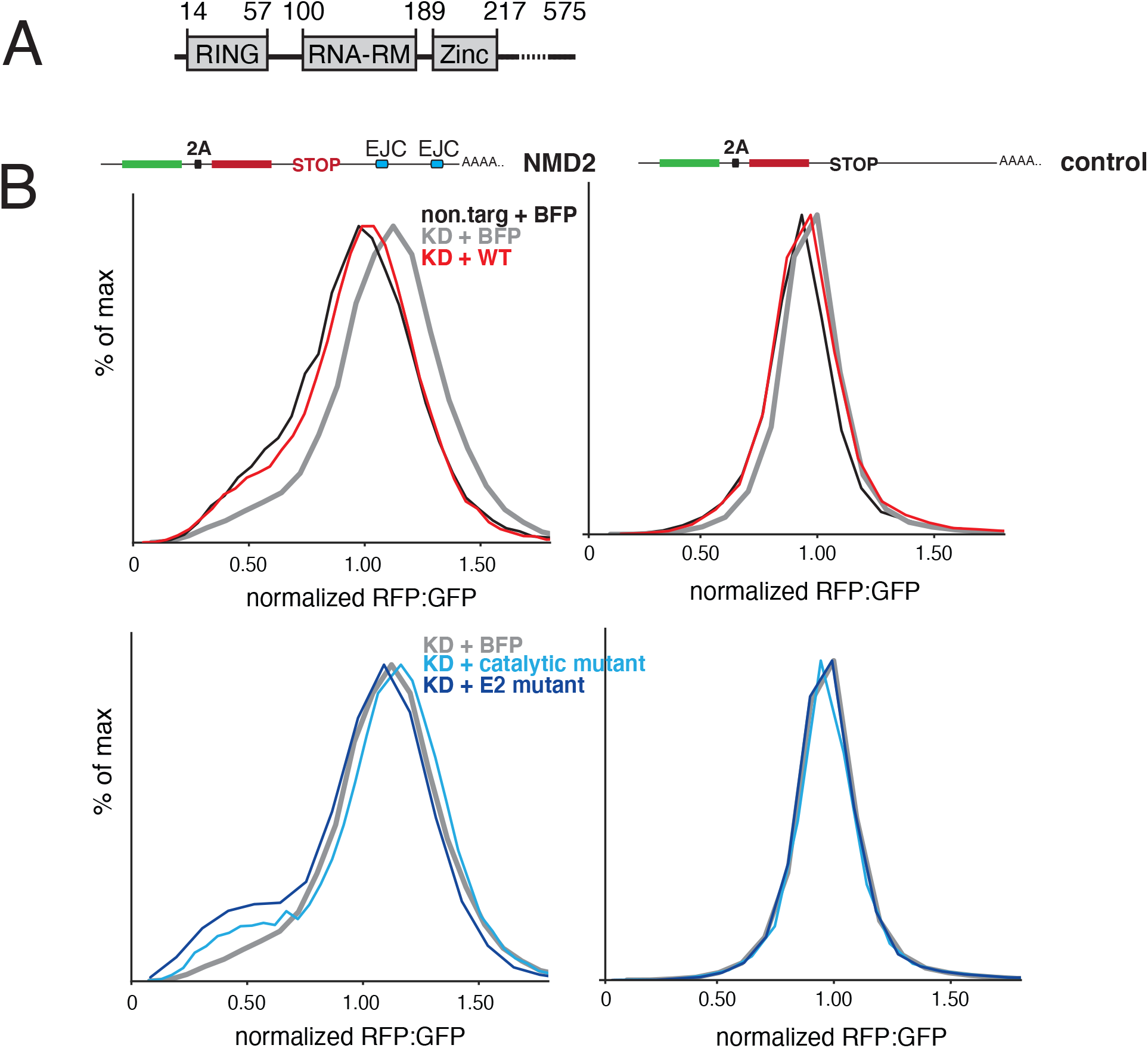
A putative role for CNOT4 in NMD-coupled protein quality control. (A) Schematic of the E3 ubiquitin ligase CNOT4 domain architecture (RNA recognition motif=RNA-RM). (B) CNOT4 depletion was rescued by expression of either wild type or two CNOT4 mutants: (i) a catalytic mutant (C14A, C17A), predicted to disrupt the folding of the CNOT4 RING domain; or (ii) an E2 mutant (L16A, C17A, C33R) that disrupts binding of CNOT4 to its cognate E2 (Albert et al., 2002). BFP was co-expressed from the rescue plasmids to allow identification of cells expressing the CNOT4 wt or mutant proteins. Displayed is a histogram of RFP:GFP fluorescence of the NMD2 and control reporters in the indicated conditions in comparison to a mock control expressing BFP alone. For comparison, data for KD+BFP is displayed in both the top and bottom histograms.

To validate the results of the screens, we first used small interfering RNA (siRNA) to deplete CNOT4, and observed a modest, but reproducible increase in the RFP:GFP ratio of our NMD reporter (Fig. 6B). Exogenous expression of wildtype CNOT4 specifically rescued the RFP:GFP ratio of the NMD reporter but had no effect on a matched control (Fig. 6B, sFig. 4C), excluding off-target effects. We then tested whether the rescue was dependent on the E3 ligase activity of CNOT4. For this, we generated two mutant constructs: (i) a mutation to the catalytic residues of the CNOT4 RING ligase based on sequence alignments with other RING-containing E3 ligases; and (ii) a mutant that disrupts binding between CNOT4 and its cognate E3 enzyme (Albert et al., 2002). Neither the catalytic mutant nor the E2 mutant were able to rescue the RFP:GFP ratio. Indeed, in the case of the E2 mutant we observed a small but reproducible dominant negative effect. To exclude off-target effects, we confirmed that depletion of CNOT4, and overexpression of the wild type or mutant protein did not affect our non-NMD control reporter. Together these data suggest that CNOT4 is specifically involved in degradation of nascent NMD protein products, in a manner dependent on its E3 ligase activity.

## DISCUSSION

Recognition of an NMD-substrate occurs co-translationally, necessarily resulting in the production of a nascent, potentially cytotoxic polypeptide chain. NMD typically reduces the mRNA level of its substrates between 2-50 fold, depending on the transcript and function of the resulting protein product: a reduction that may not be sufficient to maintain the proteostasis of the cell. As such, there has long been speculation as to whether NMD leverages an additional, post-translational pathway to directly target these nascent proteins for degradation (Chu et al., 2021; Kuroha et al., 2009, 2013; Pradhan et al., 2021; Udy & Bradley, 2021).

There are two plausible strategies by which protein degradation of NMD nascent chain occurs. Since many NMD substrates are truncated and thus likely to misfold, they expose hydrophobic degrons that will be recognized by general cytosolic quality control machinery. However, this type of uncoordinated clearance strategy would risk the cell’s exposure to transient dominant negative or gain-of-function activity of these truncated or aberrant proteins. In contrast, a coordinated protein quality control pathway that co-translationally initiates protein degradation prior to dissociation from the ribosome would be more consistent with other mRNA surveillance pathways. Indeed, tight coupling of quality control to biogenesis is a strategy used throughout biology to ensure robust and efficient clearance of mRNA and protein products that fail during their maturation (Rodrigo-Brenni & Hegde, 2012).

In the case of NMD, the lack of a robust in vitro reconstitution system; the difficulty of deconvoluting post-transcriptional versus post-translational effects on expression of NMD substrates; and the putative contribution of generalized quality control in turnover of the classical truncated NMD substrates has made it difficult to definitively identify this type of coordinated pathway. Using a fluorescent reporter strategy that addresses several of these technical challenges, we demonstrated that in mammals, NMD relies on a coupled protein quality control branch to concomitantly target the nascent protein for degradation via the ubiquitin proteasome pathway.

### A coupled protein quality control branch of NMD

We propose the following working model for protein quality control during NMD in mammals (Fig. 7). As the ribosome reaches the stop codon during translational elongation, the protein composition of the downstream mRNA serves as the primary cue for initiating NMD. At this point, the nascent polypeptide remains tethered to the ribosome via the peptidyl tRNA. We postulate that the early recognition steps between the mRNA and protein quality control branches of NMD are shared, and rely on core NMD factors such as UPF1, UPF2, UPF3b, and CASC3. We postulate that NMD-coupled quality control is thus initiated through the canonical pathway for recognition of PTC-containing mRNAs that involves binding between the ribosome, NMD factors, and the downstream EJC (Gerbracht et al., 2020; Chamieh et al., 2008; Czaplinski et al., 1998; Kim et al., 2001; le Hir et al., 2001). However, because our screens were designed to specifically query factors required for NMD-coupled protein quality control, we find substantial differences between hits identified here and those reported from earlier NMD RNA-degradation screens (Alexandrov et al., 2017; Baird et al., 2018; Sun et al., 2011; Zinshteyn et al., 2021). This discrepancy suggests that following recognition of an NMD substrate, the mRNA and protein quality control pathways diverge, relying on distinct sets of factors to target and degrade either the mRNA or nascent protein.

**Figure 7.**
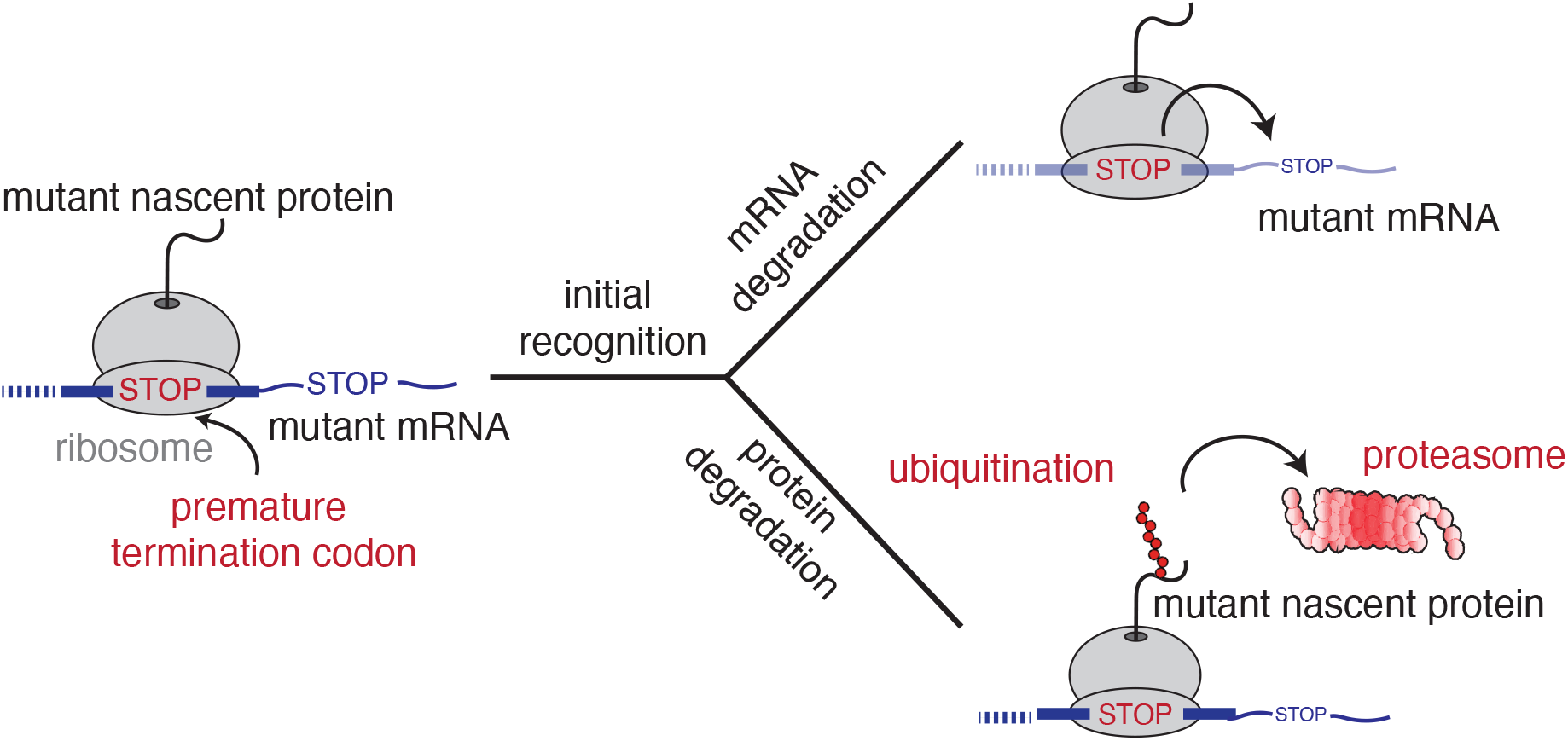
Model for NMD-coupled protein quality control. When the ribosome reaches the stop codon, NMD substrates are recognized in a context-dependent manner. These early recognition steps initiate two parallel pathways that rely on distinct suites of factors to concomitantly degrade the mRNA and nascent protein. We postulate that NMD-coupled quality control results in ubiquitination of the nascent protein prior to its release from the ribosome where it subsequently degraded by the proteasome.

We favor a model in which degradation of the nascent polypeptide is initiated prior to its release from the ribosome, as is common to other mRNA surveillance pathways and would minimize potential exposure of an aberrant protein to the cytosol. Consistent with this model we (i) found that only the nascent polypeptide tethered to the ribosome at the stop codon is subjected to NMD-coupled degradation (Fig. 1D, Fig. 2B); and (ii) we observe an NMD-specific destabilization of an intact, folded protein compared to a matched control. We therefore concluded that the nascent protein must be somehow ‘marked’ for degradation prior to its dissociation from the ribosome.

Following ubiquitination of the nascent protein, it can then be safely released into the cytosol for degradation by the proteasome. In contrast to non-stop and no-go mRNA decay where the primary cue for protein quality control is ribosome stalling (Brandman & Hegde, 2016), NMD is initiated at a stop codon and thus may utilize the typical strategy for nascent protein release and ribosome recycling. Because termination at PTCs occurs more slowly than at a canonical stop codon (Amrani et al. 2004), this additional window may be critical to allow tagging of the nascent chain for degradation prior to its release from the ribosome. However, we cannot formally exclude the possibility that this occurs simultaneously or immediately following translational termination, but prior to dissociation of the nascent chain.

### A potential role for the RQC pathway in NMD-coupled protein quality control

Several non-mutually exclusive models have been proposed for how to coordinate ubiquitination of the nascent protein chain prior to release. Experiments in *Drosophila* and *C. elegans* have suggested that at least in some systems, NMD and non-stop decay may be coupled, and levels of some mRNAs and their associated protein products are regulated by both pathways (Arribere & Fire, 2018; Hashimoto et al., 2017). A forward genetic screen in *C. elegans* further identified the canonical RQC factor Pelo (the functional ortholog of dom34/Pelota) as required for repression of an NMD reporter. Based on these and other experiments, the authors proposed a model whereby quality control by NMD in initiated by endonucleolytic cleavage of the mRNA upstream of the stop codon by SMG6. Translation of the resulting truncated mRNA would result in stalling of subsequent ribosomes at its 3’ end, triggering further repression at both the mRNA and protein level by the non-stop decay and RQC pathways (Arribere & Fire, 2018).

If a similar mechanism was occurring in mammalian cells, post-translational degradation of NMD substrates would depend on the canonical RQC factors including the E3 ubiquitin ligase LTN1, and the ribosome rescue factors pelota and HBS1. However, the majority of RQC factors were not significant hits in either of our screens, though were identified in an earlier non-stop decay screen performed using matched conditions (Hickey et al., 2020). Further, depletion of LTN1 directly did not affect our NMD reporter under conditions that robustly stabilized a non-stop decay substrate (Fig. 4C). These results suggest that at least for the class of NMD substrates represented by our reporter, NMD-coupled protein degradation does not rely on the canonical RQC pathway. Together these data suggest a functional separation of nonsense and non-stop decay in mammals, as was observed in *S. cerevisiae* (Arribere & Fire, 2018) and is consistent with the distinct molecular players identified by NMD versus non-stop mRNA decay screens (e.g. Hodgkin et al., 1989; Leeds et al., 1991; Pulak & Anderson, 1993; Wilson et al., 2007).

### Direct ubiquitination of the nascent NMD polypeptide

The simpler model for NMD-coupled protein degradation is the direct recruitment of an E3 ligase that ubiquitinates the nascent chain while it remains tethered to the ribosome. Earlier studies have suggested that UPF1, a RING domain E3 ubiquitin ligase and core NMD factor that interacts with both the ribosome and eukaryotic release factors, could carry out this role. UPF1 knockdown has been shown to stabilize protein products produced from NMD substrates mRNAs (Kuroha, Tatematsu, and Inada 2009; Kuroha et al. 2013; Feng, Jagannathan, and Bradley 2017; Park et al. 2020; Kadlec et al. 2006; Takahashi et al. 2008). Consistent with these reports, UPF1 was identified in our knockdown screen, and depletion of UPF1 stabilized both the mRNA and protein levels of our NMD reporter. However, we found that point mutations to UPF1 that specifically affect its ability to recruit its E2 ubiquitin-conjugating enzyme while leaving its ribosome-binding and helicase domains intact, did not have any effect on the protein-degradation phenotype of our reporter. We therefore concluded that UPF1 is required for NMD-coupled protein quality control, but plays a role that does not depend on its E3 ubiquitin ligase activity. To reconcile these results with previous studies, we propose that UPF1 is involved in the early recognition steps of NMD substrates, which affects both the mRNA and protein degradation branches of NMD. However, our data are inconsistent with a direct role for UPF1 in ubiquitination of the nascent polypeptide.

### A potential role for the E3 ubiquitin ligase CNOT4 in NMD-coupled protein quality control

One of the most striking hits in both our knockdown and knockout screen was the E3 ubiquitin ligase CNOT4. CNOT4 is a component of the CCR4-NOT complex, a conserved multi-subunit complex that plays a broad role in gene regulation primarily through its deadenylase activity. In NMD, the CCR4-NOT complex is recruited to transcripts through interactions between SMG7 and the CCR4-NOT subunit POP2, where it promotes deadenylation and the subsequent 3’-5’ degradation of the mRNA (Loh et al., 2013). CNOT4 is found in all eukaryotes, but is not a core structural component of the complex: in human cells it is known to cycle on and off, and its depletion does not destabilize other complex components (Jeske et al. 2006; Lau et al. 2009). CNOT4 was not identified as a significant hit in earlier screens querying NMD-mRNA levels (Alexandrov et al., 2017; Baird et al., 2018; Sun et al., 2011; Zinshteyn et al., 2021), suggesting its function is specific to the protein degradation branch of NMD.

Consistent with this model, we find that depletion of CNOT4 increases the RFP:GFP ratio of our NMD reporter by preferentially stabilizing the RFP levels, suggesting it does not markedly effect mRNA transcript levels. Though knockdown of CNOT4 had a reproducible effect in multiple cell types, the phenotype of its depletion on our NMD reporter is modest. This may be due to several contributing factors. For example, it is clear that most protein quality control pathways are highly redundant, making it challenging to observe large effects as a result of a single genetic perturbation (Rodrigo-Brenni & Hegde, 2012; Zavodszky et al., 2021). Indeed, when we generated a full null mutant, we observed compensation for loss of CNOT4, suggesting that there may be other E3 ubiquitin ligases that may be at least partially redundant. The phenotypes observed in our acute knockdown experiments are in-line with those previously reported in other systems. Because our model suggests CNOT4 may act catalytically in NMD-coupled protein quality control, very efficient depletion may be required to observe marked phenotypes.

However, the modest, but reproducible effect of CNOT4 depletion is highly specific to NMD substrates. Despite its reported role in proteasome maturation and assembly (Panasenko & Collart, 2011), CNOT4 depletion does not affect the fluorescence of our matched non-NMD reporter. We therefore concluded that CNOT4 plays a specific role in NMD-coupled protein degradation which cannot be explained by global changes to protein turnover rates. The role of CNOT4 in NMD-coupled protein degradation further appears to be distinct from its role in the CCR4-NOT complex, because CNOT1 was not a significant hit in our NMD screen, though was identified in an earlier matched CRISPRi screen for non-stop decay (Hickey et al., 2020). Finally, we demonstrate that both the E3 ligase activity of CNOT4, and its ability to bind to cognate E2 conjugating enzymes, is required for its role in NMD-coupled nascent protein degradation.

The domain architecture of CNOT4 would be consistent with a putative role in ribosome binding, given the presence of both a conserved RNA binding and zinc finger domains. Further, earlier work has implicated CNOT4 in protein quality control of non-stop mRNA decay substrates, as Not4p (the yeast ortholog of CNOT4) knockout, but not depletion of other CCR4-NOT components, stabilized truncated proteins produced from non-stop mRNAs (Dimitrova et al., 2009). The mechanistic role of CNOT4 in protein quality control of non-stop and NMD substrates in diverse eukaryotic systems thus represents an important area for future study.

### Implications of nascent protein degradation in proteostasis

The identification of a tightly coupled protein degradation branch of NMD has several immediate implications. Most notably, destabilization at the post-translational level will increase the suppression of NMD substrates. Though we find the effects of NMD-coupled protein degradation on our reporters to be modest (∼2-fold), in the context of the cell or an organism, this additional level of regulation may be critical to prevent deleterious or off-target effects. Effects on these fluorescent reporters, which are both over-expressed and in which phenotypes require degradation of the remarkably stable RFP moiety, may also underestimate the true effect size on an endogenous substrate.

There are numerous physiologically relevant examples where NMD’s role in transcriptome regulation, and subsequent production of potentially aberrant proteins, require stringent clearance of the nascent product. During histone production, synthesis must be tightly regulated in a manner coupled to the progression of the cell cycle, and the production of even small amounts of down-regulated proteins could be problematic. Our results also have implications for viral infection. Co-translational protein degradation is thought to be a key source of peptides for MHC presentation (Balistreri et al., 2014; Fontaine et al., 2018; Wada et al., 2018; Yewdell & Nicchitta, 2006), with viral messages often targeted by NMD (Balistreri et al., 2014; Fontaine et al., 2018; Wada et al., 2018). Factors such as CNOT4 could mediate this process, promoting the immunological presentation of these peptides.

Finally, NMD plays an important role in a wide range of genetic diseases: over one third of all human genetic disorders are caused by PTC-creating mutations, including muscular dystrophy and cystic fibrosis. While generally protective, for numerous disease-causing mutations the NMD pathway contributes to pathogenesis by suppressing expression of partially functional mutant proteins (∼11% of mutations that cause human disease (Mort et al., 2008)). The characterization of a second, parallel branch of NMD and the factors involved in NMD-coupled protein quality control therefore represent novel targets for the therapeutic treatment of human disease.

## ACKNOWLEDGEMENTS

We thank Tino Pleiner, Joseph Replogle and the Voorhees lab for discussion and Simon Byrne and Patrick Smyth for help with data processing. Flow cytometry experiments were performed at the Caltech flow cytometry facility. The Whitehead Institute Genome Technology core performed sequencing for genome-wide screens. This work was supported by the Heritage Medical Research Institute, the Kinship Foundation, the Pew-Stewart Foundation and by the Millard and Muriel Jacobs Genetics and Genomics Laboratory at California Institute of Technology. A.J.I. was supported by a Caltech BBE postdoctoral fellowship and a grant from the Larry L. Hillblom Foundation and A.G. by a Human Frontier Science Program fellowship. The three co-first authors are joint authors and as such they can prioritize their names when adding this paper to their resumes.

## Supplementary Materials

Materials and Methods

Figs. S1-S4

Supplementary Tables 1-3

## REFERENCES

Albert, T. K., Hanzawa, H., Legtenberg, Y. I. A., de Ruwe, M. J., van den Heuvel, F. A. J., Collart, M. A., Boelens, R., & Timmers, H. T. M. (2002). Identification of a ubiquitin--protein ligase subunit within the CCR4--NOT transcription repressor complex. The EMBO Journal, 21(3), 355–364.

Alexandrov, A., Shu, M.-D., & Steitz, J. A. (2017). Fluorescence amplification method for forward genetic discovery of factors in human mRNA degradation. Molecular Cell, 65(1), 191–201.

Amrani, N., Ganesan, R., Kervestin, S., Mangus, D. A., Ghosh, S., & Jacobson, A. (2004). A faux 3′-UTR promotes aberrant termination and triggers nonsense-mediated mRNA decay. Nature, 432(7013), 112–118.

Anczuków, O., Ware, M. D., Buisson, M., Zetoune, A. B., Stoppa-Lyonnet, D., Sinilnikova, O. M., & Mazoyer, S. (2008). Does the nonsense-mediated mRNA decay mechanism prevent the synthesis of truncated BRCA1, CHK2, and p53 proteins? Human Mutation, 29(1), 65–73.

Arribere, J. A., & Fire, A. Z. (2018). Nonsense mRNA suppression via nonstop decay. Elife, 7, e33292.

Baird, T. D., Cheng, K. C.-C., Chen, Y.-C., Buehler, E., Martin, S. E., Inglese, J., & Hogg, J. R. (2018). ICE1 promotes the link between splicing and nonsense-mediated mRNA decay. Elife, 7, e33178.

Balistreri, G., Horvath, P., Schweingruber, C., Zünd, D., McInerney, G., Merits, A., Mühlemann, O., Azzalin, C., & Helenius, A. (2014). The host nonsense-mediated mRNA decay pathway restricts mammalian RNA virus replication. Cell Host \& Microbe, 16(3), 403–411.

Balleza, E., Kim, J. M., & Cluzel, P. (2018). Systematic characterization of maturation time of fluorescent proteins in living cells. Nature Methods, 15(1), 47–51.

Ballut, L., Marchadier, B., Baguet, A., Tomasetto, C., Séraphin, B., & le Hir, H. (2005). The exon junction core complex is locked onto RNA by inhibition of eIF4AIII ATPase activity. Nature Structural \& Molecular Biology, 12(10), 861–869.

Becker, T., Armache, J.-P., Jarasch, A., Anger, A. M., Villa, E., Sieber, H., Motaal, B. A., Mielke, T., Berninghausen, O., & Beckmann, R. (2011). Structure of the no-go mRNA decay complex Dom34--Hbs1 bound to a stalled 80S ribosome. Nature Structural \& Molecular Biology, 18(6), 715–720.

Behm-Ansmant, I., Gatfield, D., Rehwinkel, J., Hilgers, V., & Izaurralde, E. (2007). A conserved role for cytoplasmic poly (A)-binding protein 1 (PABPC1) in nonsense-mediated mRNA decay. The EMBO Journal, 26(6), 1591–1601.

Belgrader, P., Cheng, J., & Maquat, L. E. (1993). Evidence to implicate translation by ribosomes in the mechanism by which nonsense codons reduce the nuclear level of human triosephosphate isomerase mRNA. Proceedings of the National Academy of Sciences, 90(2), 482–486.

Brandman, O., & Hegde, R. S. (2016). Ribosome-associated protein quality control. Nature Structural & Molecular Biology, 23(1), 7–15. https://doi.org/10.1038/nsmb.3147

Brandman, O., Stewart-Ornstein, J., Wong, D., Larson, A., Williams, C. C., Li, G.-W., Zhou, S., King, D., Shen, P. S., Weibezahn, J., & others. (2012). A ribosome-bound quality control complex triggers degradation of nascent peptides and signals translation stress. Cell, 151(5), 1042–1054.

Bruce, S. R., & Wilkinson, M. F. (2003). Nonsense-mediated decay: A surveillance pathway that detects faulty TCR and BCR transcripts. Research Signpost, Trivandrum, India.

Celik, A., Baker, R., He, F., & Jacobson, A. (2017). High-resolution profiling of NMD targets in yeast reveals translational fidelity as a basis for substrate selection. Rna, 23(5), 735–748.

Chamieh, H., Ballut, L., Bonneau, F., & le Hir, H. (2008). NMD factors UPF2 and UPF3 bridge UPF1 to the exon junction complex and stimulate its RNA helicase activity. Nature Structural \& Molecular Biology, 15(1), 85–93.

Chang, J. C., & Kan, Y. W. (1979). Β0 Thalassemia, a Nonsense Mutation in Man. Proceedings of the National Academy of Sciences of the United States of America, 76(6), 2886–2889. https://doi.org/10.1073/pnas.76.6.2886

Chen, C.-Y. A., & Shyu, A.-B. (2003). Rapid deadenylation triggered by a nonsense codon precedes decay of the RNA body in a mammalian cytoplasmic nonsense-mediated decay pathway. Molecular and Cellular Biology, 23(14), 4805–4813.

Cho, H., Kim, K. M., & Kim, Y. K. (2009). Human proline-rich nuclear receptor coregulatory protein 2 mediates an interaction between mRNA surveillance machinery and decapping complex. Molecular Cell, 33(1), 75–86.

Choe, J., Ahn, S. H., & Kim, Y. K. (2014). The mRNP remodeling mediated by UPF1 promotes rapid degradation of replication-dependent histone mRNA. Nucleic Acids Research, 42(14), 9334–9349.

Chu, V., Feng, Q., Lim, Y., & Shao, S. (2021). Selective destabilization of polypeptides synthesized from NMD-targeted transcripts. Molecular Biology of the Cell, mbc--E21.

Collart, M. A. (2016). The Ccr4-Not complex is a key regulator of eukaryotic gene expression. Wiley Interdisciplinary Reviews: RNA, 7(4), 438–454.

Czaplinski, K., Ruiz-Echevarria, M. J., Paushkin, S. v, Han, X., Weng, Y., Perlick, H. A., Dietz, H. C., Ter-Avanesyan, M. D., & Peltz, S. W. (1998). The surveillance complex interacts with the translation release factors to enhance termination and degrade aberrant mRNAs. Genes \& Development, 12(11), 1665–1677.

Defenouillère, Q., Yao, Y., Mouaikel, J., Namane, A., Galopier, A., Decourty, L., Doyen, A., Malabat, C., Saveanu, C., Jacquier, A., & others. (2013). Cdc48-associated complex bound to 60S particles is required for the clearance of aberrant translation products. Proceedings of the National Academy of Sciences, 110(13), 5046–5051.

Dietz, H. C., Valle, D., Francomano, C. A., Kendzior, R. J., Pyeritz, R. E., & Cutting, G. R. (1993). The skipping of constitutive exons in vivo induced by nonsense mutations. Science, 259(5095), 680–683.

Dimitrova, L. N., Kuroha, K., Tatematsu, T., & Inada, T. (2009). Nascent peptide-dependent translation arrest leads to Not4p-mediated protein degradation by the proteasome. Journal of Biological Chemistry, 284(16), 10343–10352.

Doma, M. K., & Parker, R. (2006). Endonucleolytic cleavage of eukaryotic mRNAs with stalls in translation elongation. Nature, 440(7083), 561–564.

Dostie, J., & Dreyfuss, G. (2002). Translation is required to remove Y14 from mRNAs in the cytoplasm. Current Biology, 12(13), 1060–1067.

Eberle, A. B., Lykke-Andersen, S., Mühlemann, O., & Jensen, T. H. (2009). SMG6 promotes endonucleolytic cleavage of nonsense mRNA in human cells. Nature Structural \& Molecular Biology, 16(1), 49–55.

Feng, Q., Jagannathan, S., & Bradley, R. K. (2017). The RNA Surveillance Factor UPF1 Represses Myogenesis via Its E3 Ubiquitin Ligase Activity. Molecular Cell, 67(2), 239–251.e6. https://doi.org/10.1016/j.molcel.2017.05.034

Fontaine, K. A., Leon, K. E., Khalid, M. M., Tomar, S., Jimenez-Morales, D., Dunlap, M., Kaye, J. A., Shah, P. S., Finkbeiner, S., Krogan, N. J., & others. (2018). The cellular NMD pathway restricts Zika virus infection and is targeted by the viral capsid protein. MBio, 9(6), e02126.–18.

Frischmeyer, P. A., van Hoof, A., O’Donnell, K., Guerrerio, A. L., Parker, R., & Dietz, H. C. (2002). An mRNA surveillance mechanism that eliminates transcripts lacking termination codons. Science, 295(5563), 2258–2261.

Gehring, N. H., Neu-Yilik, G., Schell, T., Hentze, M. W., & Kulozik, A. E. (2003). Y14 and hUpf3b form an NMD-activating complex. Molecular Cell, 11(4), 939–949.

Gerbracht, J. v, Boehm, V., Britto-Borges, T., Kallabis, S., Wiederstein, J. L., Ciriello, S., Aschemeier, D. U., Krüger, M., Frese, C. K., Altmüller, J., & others. (2020). CASC3 promotes transcriptome-wide activation of nonsense-mediated decay by the exon junction complex. Nucleic Acids Research, 48(15), 8626–8644.

Gilbert, L. A., Horlbeck, M. A., Adamson, B., Villalta, J. E., Chen, Y., Whitehead, E. H., Guimaraes, C., Panning, B., Ploegh, H. L., Bassik, M. C., & others. (2014). Genome-scale CRISPR-mediated control of gene repression and activation. Cell, 159(3), 647–661.

Hall, G. W., & Thein, S. (1994). Nonsense codon mutations in the terminal exon of the beta-globin gene are not associated with a reduction in beta-mRNA accumulation: a mechanism for the phenotype of dominant beta-thalassemia.

Hanzawa, H., de Ruwe, M. J., Albert, T. K., van der Vliet, P. C., Timmers, H. T. M., & Boelens, R. (2001). The Structure of the C4C4RING finger of human NOT4 reveals features distinct from those of C3HC4 RING fingers. Journal of Biological Chemistry, 276(13), 10185–10190.

Hart, T., Tong, A. H. Y., Chan, K., van Leeuwen, J., Seetharaman, A., Aregger, M., Chandrashekhar, M., Hustedt, N., Seth, S., Noonan, A., Habsid, A., Sizova, O., Nedyalkova, L., Climie, R., Tworzyanski, L., Lawson, K., Sartori, M. A., Alibeh, S., Tieu, D., … Moffat, J. (2017). Evaluation and Design of Genome-Wide CRISPR/SpCas9 Knockout Screens. G3: Genes, Genomes, Genetics, 7(8), 2719–2727. https://doi.org/10.1534/g3.117.041277

Hashimoto, Y., Takahashi, M., Sakota, E., & Nakamura, Y. (2017). Nonstop-mRNA decay machinery is involved in the clearance of mRNA 5′-fragments produced by RNAi and NMD in Drosophila melanogaster cells. Biochemical and Biophysical Research Communications, 484(1), 1–7.

He, F., Li, X., Spatrick, P., Casillo, R., Dong, S., & Jacobson, A. (2003). Genome-Wide Analysis of mRNAs Regulated by the Nonsense-Mediated and 5′ to 3′ mRNA Decay Pathways in Yeast. Molecular Cell, 12(6), 1439–1452. https://doi.org/https://doi.org/10.1016/S1097-2765(03)00446-5

Hickey, K. L., Dickson, K., Cogan, J. Z., Replogle, J. M., Schoof, M., D’Orazio, K. N., Sinha, N. K., Hussmann, J. A., Jost, M., Frost, A., & others. (2020). GIGYF2 and 4EHP inhibit translation initiation of defective messenger RNAs to assist ribosome-associated quality control. Molecular Cell, 79(6), 950–962.

Hodgkin, J., Papp, A., Pulak, R., Ambros, V., & Anderson, P. (1989). A new kind of informational suppression in the nematode Caenorhabditis elegans. Genetics, 123(2), 301–313.

Hoek, T. A., Khuperkar, D., Lindeboom, R. G. H., Sonneveld, S., Verhagen, B. M. P., Boersma, S., Vermeulen, M., & Tanenbaum, M. E. (2019). Single-molecule imaging uncovers rules governing nonsense-mediated mRNA decay. Molecular Cell, 75(2), 324–339.

Horlbeck, M. A., Gilbert, L. A., Villalta, J. E., Adamson, B., Pak, R. A., Chen, Y., Fields, A. P., Park, C. Y., Corn, J. E., Kampmann, M., & others. (2016). Compact and highly active nextgeneration libraries for CRISPR-mediated gene repression and activation. Elife, 5, e19760.

Hoskins, A. A., & Moore, M. J. (2012). The spliceosome: a flexible, reversible macromolecular machine. Trends in Biochemical Sciences, 37(5), 179–188.

Huntzinger, E., Kashima, I., Fauser, M., Saulière, J., & Izaurralde, E. (2008). SMG6 is the catalytic endonuclease that cleaves mRNAs containing nonsense codons in metazoan. Rna, 14(12), 2609–2617.

Jeske, M., Meyer, S., Temme, C., Freudenreich, D., & Wahle, E. (2006). Rapid ATP-dependent deadenylation of nanos mRNA in a cell-free system from Drosophila embryos. Journal of Biological Chemistry, 281(35), 25124–25133.

Jing, Z., Sun, X., Qian, Y., & Maquiat, L. E. (1998). Intron function in the nonsense-mediated decay of β-globin mRNA: Indications that pre-mRNA splicing in the nucleus can influence mRNA translation in the cytoplasm. RNA, 4(7), 801–815. https://doi.org/10.1017/S1355838298971849

Jost, M., Chen, Y., Gilbert, L. A., Horlbeck, M. A., Krenning, L., Menchon, G., Rai, A., Cho, M. Y., Stern, J. J., Prota, A. E., & others. (2017). Combined CRISPRi/a-based chemical genetic screens reveal that rigosertib is a microtubule-destabilizing agent. Molecular Cell, 68(1), 210–223.

Juszkiewicz, S., Chandrasekaran, V., Lin, Z., Kraatz, S., Ramakrishnan, V., & Hegde, R. S. (2018). ZNF598 is a quality control sensor of collided ribosomes. Molecular Cell, 72(3), 469–481.

Karam, R., Carvalho, J., Bruno, I., Graziadio, C., Senz, J., Huntsman, D., Carneiro, F., Seruca, R., Wilkinson, M. F., & Oliveira, C. (2008). The NMD mRNA surveillance pathway downregulates aberrant E-cadherin transcripts in gastric cancer cells and in CDH1 mutation carriers. Oncogene, 27(30), 4255–4260. https://doi.org/10.1038/onc.2008.62

Kerr, T. P., Sewry, C. A., Robb, S. A., & Roberts, R. G. (2001). Long mutant dystrophins and variable phenotypes: evasion of nonsense-mediated decay? Human Genetics, 109(4), 402–407.

Kim, V. N., Kataoka, N., & Dreyfuss, G. (2001). Role of the nonsense-mediated decay factor hUpf3 in the splicing-dependent exon-exon junction complex. Science, 293(5536), 1832–1836.

Kugler, W., Enssle, J., Hentze, M. W., & Kulozik, A. E. (1995). Nuclear degradation of nonsense mutated $β$-globin mRNA: a post-transcriptional mechanism to protect heterozygotes from severe clinical manifestations of $β$-thalassemia? Nucleic Acids Research, 23(3), 413–418.

Kuroha, K., Ando, K., Nakagawa, R., & Inada, T. (2013). The Upf factor complex interacts with aberrant products derived from mRNAs containing a premature termination codon and facilitates their proteasomal degradation. Journal of Biological Chemistry, 288(40), 28630–28640.

Kuroha, K., Tatematsu, T., & Inada, T. (2009). Upf1 stimulates degradation of the product derived from aberrant messenger RNA containing a specific nonsense mutation by the proteasome. EMBO Reports, 10(11), 1265–1271.

Lai, T., Cho, H., Liu, Z., Bowler, M. W., Piao, S., Parker, R., Kim, Y. K., & Song, H. (2012). Structural basis of the PNRC2-mediated link between mRNA surveillance and decapping. Structure, 20(12), 2025–2037.

Lau, N.-C., Kolkman, A., van Schaik, F. M. A., Mulder, K. W., Pijnappel, W. W. M. P., Heck, A. J. R., & Timmers, H. T. M. (2009). Human Ccr4--Not complexes contain variable deadenylase subunits. Biochemical Journal, 422(3), 443–453.

le Hir, H., Gatfield, D., Izaurralde, E., & Moore, M. J. (2001). The exon--exon junction complex provides a binding platform for factors involved in mRNA export and nonsense-mediated mRNA decay. The EMBO Journal, 20(17), 4987–4997.

le Hir, H., Izaurralde, E., Maquat, L. E., & Moore, M. J. (2000). The spliceosome deposits multiple proteins 20--24 nucleotides upstream of mRNA exon--exon junctions. The EMBO Journal, 19(24), 6860–6869.

Leeds, P., Peltz, S. W., Jacobson, A., & Culbertson, M. R. (1991). The product of the yeast UPF1 gene is required for rapid turnover of mRNAs containing a premature translational termination codon. Genes \& Development, 5(12a), 2303–2314.

Lelivelt, M. J., & Culbertson, M. R. (1999). Yeast Upf Proteins Required for RNA Surveillance Affect Global Expression of the Yeast Transcriptome. Molecular and Cellular Biology, 19(10), 6710–6719. https://doi.org/10.1128/MCB.19.10.6710

Li, W., Xu, H., Xiao, T., Cong, L., Love, M. I., Zhang, F., Irizarry, R. A., Liu, J. S., Brown, M., & Liu, X. S. (2014). MAGeCK enables robust identification of essential genes from genomescale CRISPR/Cas9 knockout screens. Genome Biology, 15(12), 1–12.

Loh, B., Jonas, S., & Izaurralde, E. (2013). The SMG5--SMG7 heterodimer directly recruits the CCR4--NOT deadenylase complex to mRNAs containing nonsense codons via interaction with POP2. Genes \& Development, 27(19), 2125–2138.

Losson, R., & Lacroute, F. (1979). Interference of nonsense mutations with eukaryotic messenger RNA stability. Proceedings of the National Academy of Sciences, 76(10), 5134–5137.

Lykke-Andersen, J., Shu, M. di, & Steitz, J. A. (2000). Human Upf proteins target an mRNA for nonsense-mediated decay when downstream of a termination codon. Cell, 103(7), 1121–1131. https://doi.org/10.1016/S0092-8674(00)00214-2

Lyumkis, D., dos Passos, D. O., Tahara, E. B., Webb, K., Bennett, E. J., Vinterbo, S., Potter, C. S., Carragher, B., & Joazeiro, C. A. P. (2014). Structural basis for translational surveillance by the large ribosomal subunit-associated protein quality control complex. Proceedings of the National Academy of Sciences, 111(45), 15981–15986.

Maquat, L. E., Kinniburgh, A. J., Rachmilewitz, E. A., & Ross, J. (1981). Unstable beta-globin mRNA in mRNA-deficient beta o thalassemia. Cell, 27(3 Pt 2), 543–553. http://www.ncbi.nlm.nih.gov/pubmed/6101206

Marino, J., von Heijne, G., & Beckmann, R. (2016). Small protein domains fold inside the ribosome exit tunnel. FEBS Letters, 590(5), 655–660.

Mendell, J. T., Sharifi, N. A., Meyers, J. L., Martinez-Murillo, F., & Dietz, H. C. (2004). Nonsense surveillance regulates expression of diverse classes of mammalian transcripts and mutes genomic noise. Nature Genetics, 36(10), 1073–1078. https://doi.org/10.1038/ng1429

Mitchell, P., & Tollervey, D. (2003). An NMD pathway in yeast involving accelerated deadenylation and exosome-mediated 3′→ 5′ degradation. Molecular Cell, 11(5), 1405–1413.

Mort, M., Ivanov, D., Cooper, D. N., & Chuzhanova, N. A. (2008). A meta-analysis of nonsense mutations causing human genetic disease. Human Mutation, 29(8), 1037–1047.

Nagy, E., & Maquat, L. E. (1998). A rule for termination-codon position within intron-containing genes: when nonsense affects RNA abundance. Trends in Biochemical Sciences, 23(6), 198–199. https://doi.org/https://doi.org/10.1016/S0968-0004(98)01208-0

Nott, A., le Hir, H., & Moore, M. J. (2004). Splicing enhances translation in mammalian cells: an additional function of the exon junction complex. Genes \& Development, 18(2), 210–222.

O’Sullivan, B. P. (2014). Targeting nonsense-mediated cystic fibrosis: is it premature to stop now? The Lancet Respiratory Medicine, 2(7), 509–511. https://doi.org/10.1016/S2213-2600(14)70108-0

Palacios, I. M., Gatfield, D., St Johnston, D., & Izaurralde, E. (2004). An eIF4AIII-containing complex required for mRNA localization and nonsense-mediated mRNA decay. Nature, 427(6976), 753–757.

Panasenko, O. O., & Collart, M. A. (2011). Not4 E3 ligase contributes to proteasome assembly and functional integrity in part through Ecm29. Molecular and Cellular Biology, 31(8), 1610–1623.

Pereverzev, A. P., Gurskaya, N. G., Ermakova, G. v, Kudryavtseva, E. I., Markina, N. M., Kotlobay, A. A., Lukyanov, S. A., Zaraisky, A. G., & Lukyanov, K. A. (2015). Method for quantitative analysis of nonsense-mediated mRNA decay at the single cell level. Scientific Reports, 5(1), 1–10.

Perrin-Vidoz, L., Sinilnikova, O. M., Stoppa-Lyonnet, D., Lenoir, G. M., & Mazoyer, S. (2002). The nonsense-mediated mRNA decay pathway triggers degradation of most BRCA1 mRNAs bearing premature termination codons. Human Molecular Genetics, 11(23), 2805–2814.

Pisareva, V. P., Skabkin, M. A., Hellen, C. U. T., Pestova, T. v, & Pisarev, A. v. (2011). Dissociation by Pelota, Hbs1 and ABCE1 of mammalian vacant 80S ribosomes and stalled elongation complexes. The EMBO Journal, 30(9), 1804–1817.

Popp, M. W.-L., & Maquat, L. E. (2013). Organizing principles of mammalian nonsense-mediated mRNA decay. Annual Review of Genetics, 47, 139–165.

Pradhan, A. K., Kandasamy, G., Chatterjee, U., Bharadwaj, A., Mathew, S. J., Dohmen, R. J., & Palanimurugan, R. (2021). Ribosome-associated quality control mediates degradation of the premature translation termination product Orf1p of ODC antizyme mRNA. FEBS Letters.

Pulak, R., & Anderson, P. (1993). mRNA surveillance by the Caenorhabditis elegans smg genes. Genes \& Development, 7(10), 1885–1897.

Ramage, H. R., Kumar, G. R., Verschueren, E., Johnson, J. R., von Dollen, J., Johnson, T., Newton, B., Shah, P., Horner, J., Krogan, N. J., & others. (2015). A combined proteomics/genomics approach links hepatitis C virus infection with nonsense-mediated mRNA decay. Molecular Cell, 57(2), 329–340.

Reddy, J. C., Morris, J. C., Wang, J., English, M. A., Haber, D. A., Shi, Y., & Licht, J. D. (1995). WT1-mediated transcriptional activation is inhibited by dominant negative mutant proteins. Journal of Biological Chemistry, 270(18), 10878–10884.

Rehwinkel, J., Letunic, I., Raes, J., Bork, P., & Izaurralde, E. (2005). Nonsense-mediated mRNA decay factors act in concert to regulate common mRNA targets. RNA (New York, N.Y.), 11(10), 1530–1544. https://doi.org/10.1261/rna.2160905

Rendón, O. Z., Fredrickson, E. K., Howard, C. J., van Vranken, J., Fogarty, S., Tolley, N. D., Kalia, R., Osuna, B. A., Shen, P. S., Hill, C. P., & others. (2018). Vms1p is a release factor for the ribosome-associated quality control complex. Nature Communications, 9(1), 1–9.

Rodrigo-Brenni, M. C., & Hegde, R. S. (2012). Design principles of protein biosynthesis-coupled quality control. Developmental Cell, 23(5), 896–907.

Rosenbluh, J., Xu, H., Harrington, W., Gill, S., Wang, X., Vazquez, F., Root, D. E., Tsherniak, A., & Hahn, W. C. (2017). Complementary information derived from CRISPR Cas9 mediated gene deletion and suppression. Nature Communications, 8(1), 1–8.

Shao, S., Brown, A., Santhanam, B., & Hegde, R. S. (2015). Structure and assembly pathway of the ribosome quality control complex. Molecular Cell, 57(3), 433–444.

Shao, S., Malsburg, K. von der, & Hegde, R. S. (2013). Listerin-Dependent Nascent Protein Ubiquitination Relies on Ribosome Subunit Dissociation. Molecular Cell, 50(5), 637–648.

Shao, S., Murray, J., Brown, A., Taunton, J., Ramakrishnan, V., & Hegde, R. S. (2016). Decoding mammalian ribosome-mRNA states by translational GTPase complexes. Cell, 167(5), 1229–1240.

Shoemaker, C. J., & Green, R. (2012). Translation drives mRNA quality control. Nature Structural & Molecular Biology, 19(6), 594–601.

Silva, A.L, Ribeiro, P., Inacio, A., Liebhaber, S.A., and Romao, L. (2008). Proximity of poly(A)-binding protein to a premature termination codon inhibits mammalian nonsense-mediated mRNA decay. RNA, 14(3), 563–576.

Singh, G., Rebbapragada, I., & Lykke-Andersen, J. (2008). A competition between stimulators and antagonists of Upf complex recruitment governs human nonsense-mediated mRNA decay. PLoS Biology, 6(4), e111. https://doi.org/10.1371/journal.pbio.0060111

Sun, Y., Yang, P., Zhang, Y., Bao, X., Li, J., Hou, W., Yao, X., Han, J., & Zhang, H. (2011). A genome-wide RNAi screen identifies genes regulating the formation of P bodies in C. elegans and their functions in NMD and RNAi. Protein \& Cell, 2(11), 918–939.

Takahashi, S., Araki, Y., Ohya, Y., Sakuno, T., Hoshino, S. I., Kontani, K., Nishina, H., & Katada, T. (2008). Upf1 potentially serves as a RING-related E3 ubiquitin ligase via its association with Upf3 in yeast. Rna, 14(9), 1950–1958. https://doi.org/10.1261/rna.536308

Takahashi, S., Araki, Y., Sakuno, T., & Katada, T. (2003). Interaction between Ski7p and Upf1p is required for nonsense-mediated 3′-to-5′ mRNA decay in yeast. The EMBO Journal, 22(15), 3951–3959.

Thein, S. L., Hesketh, C., Taylor, P., Temperley, I. J., Hutchinson, R. M., Old, J. M., Wood, W. G., Clegg, J. B., & Weatherall, D. J. (1990). Molecular basis for dominantly inherited inclusion body beta-thalassemia. Proceedings of the National Academy of Sciences, 87(10), 3924–3928.

Udy, D. B., & Bradley, R. K. (2021). Nonsense-mediated mRNA decay utilizes complementary mechanisms to suppress mRNA and protein accumulation. BioRxiv.

van Hoof, A., Frischmeyer, P. A., Dietz, H. C., & Parker, R. (2002). Exosome-mediated recognition and degradation of mRNAs lacking a termination codon. Science, 295(5563), 2262–2264.

Verma, R., Oania, R. S., Kolawa, N. J., & Deshaies, R. J. (2013). Cdc48/p97 promotes degradation of aberrant nascent polypeptides bound to the ribosome. Elife, 2, e00308.

Verma, R., Reichermeier, K. M., Burroughs, A. M., Oania, R. S., Reitsma, J. M., Aravind, L., & Deshaies, R. J. (2018). Vms1 and ANKZF1 peptidyl-tRNA hydrolases release nascent chains from stalled ribosomes. Nature, 557(7705), 446–451.

Wada, M., Lokugamage, K. G., Nakagawa, K., Narayanan, K., & Makino, S. (2018). Interplay between coronavirus, a cytoplasmic RNA virus, and nonsense-mediated mRNA decay pathway. Proceedings of the National Academy of Sciences, 115(43), E10157--E10166.

Wang, J., Vock, V. M., Li, S., Olivas, O. R., & Wilkinson, M. F. (2002). A quality control pathway that down-regulates aberrant T-cell receptor (TCR) transcripts by a mechanism requiring UPF2 and translation. Journal of Biological Chemistry, 277(21), 18489–18493.

Wang, Y., Wang, F., Wang, R., Zhao, P., & Xia, Q. (2015). 2A self-cleaving peptide-based multigene expression system in the silkworm Bombyx mori. Scientific Reports, 5(1), 1–10.

Ware, M. D., DeSilva, D., Sinilnikova, O. M., Stoppa-Lyonnet, D., Tavtigian, S. v, & Mazoyer, S. (2006). Does nonsense-mediated mRNA decay explain the ovarian cancer cluster region of the BRCA2 gene? Oncogene, 25(2), 323–328. https://doi.org/10.1038/sj.onc.1209033

Wilson, M. A., Meaux, S., & van Hoof, A. (2007). A genomic screen in yeast reveals novel aspects of nonstop mRNA metabolism. Genetics, 177(2), 773–784.

Yamashita, A., Chang, T.-C., Yamashita, Y., Zhu, W., Zhong, Z., Chen, C.-Y. A., & Shyu, A.-B. (2005). Concerted action of poly (A) nucleases and decapping enzyme in mammalian mRNA turnover. Nature Structural \& Molecular Biology, 12(12), 1054–1063.

Yewdell, J. W., & Nicchitta, C. v. (2006). The DRiP hypothesis decennial: support, controversy, refinement and extension. Trends in Immunology, 27(8), 368–373.

Zavodszky, E., Peak-Chew, S.-Y., Juszkiewicz, S., Narvaez, A. J., & Hegde, R. S. (2021). Identification of a quality-control factor that monitors failures during proteasome assembly. Science, 373(6558), 998–1004.

Zhang, J., & Maquat, L. E. (1997). Evidence that translation reinitiation abrogates nonsense-mediated mRNA decay in mammalian cells. The EMBO Journal, 16(4), 826–833.

Zinshteyn, B., Sinha, N. K., Enam, S. U., Koleske, B., & Green, R. (2021). Translational repression of NMD targets by GIGYF2 and EIF4E2. PLoS Genetics, 17(10), e1009813.

