## Supplemental information for "Coupled protein quality control during nonsense mediated mRNA decay"

### MATERIALS AND METHODS

Plasmids and antibodies

Reporter constructs for expression in mammalian cells were generated in either the pcDNA5/FRT/TO (Thermo Scientific) backbone (for expression in HEK293T cells) or the SFFV-tet3G lentiviral backbone with a 3' WPRE element (for inducible expression in K562 cells, from (Jost et al., 2017). To create the NMD reporters described in Fig. 1, a fragment of the beta-globin gene (spanning the last 221 nucleotides of exon 2 (the last 35 nucleotides for inert EJC), intron 2 and 129 nucleotides of exon 3) was amplified via PCR from human genomic DNA as described previously (Pereverzev et al., 2015). Either one or two copies were inserted into the 3' UTR of a plasmid encoding GFP-P2A-RFP to generate NMD1 and NMD2 respectively. In the lentiviral constructs, the reporter was inserted in reverse orientation to prevent splicing of the introns during lentiviral production. The presence of functional introns was checked via PCR, using primers that should span the introns (Fig. S1B). The RNeasy kit (74104, Qiagen) was used to purify total RNA from HEK293T transiently expressing the NMD reporters. cDNA was obtained by reverse transcription using the SuperScript III First Strand Synthesis SuperMix (11752, Invitrogen). PCR amplification from this cDNA generated a shorter fragment than that of the reporter plasmids, indicating the introns have been spliced out efficiently.

Control constructs were created by replacing the P2A site with a glycine-serine linker of identical length (Fig. 1D), reversing the order of the GFP and RFP (Fig. S1D), or appending the villin headpiece domain (bVHP) downstream of the RFP (Fig. S1E). A FLAG tag was appended to the N-terminus of RFP for immunoprecipitation (Fig. 2C). Note: the mCherry and mEGFP versions of the GFP and RFP were used throughout this study, but for simplicity are referred to as GFP and RFP throughout.

cDNA for UPF1 was acquired from Addgene (Cat #99146) and cloned after a BFP-P2A sequence in a lentiviral backbone with an EF1-a promoter from an upstream ubiquitious chromatin opening element (UCOE). A mutant of UPF1 with mutations in the RING domain (S134A, N148A, T149A) that disrupts binding with E2 ligases was also acquired from Addgene (Cat #99144). DNA encoding CNOT4 was ordered as gblocks from IDT. Mutations to the catalytic cysteines of the RING domain of CNOT4 (C14A, C17A) were chosen by alignment to other E3 ligase family members. Mutations to prevent CNOT4 from binding to E2 proteins (L16A, C17A, C33R) were taken from (Albert et al., 2002).

The following antibodies were used in this study: FLAG (A2220, Sigma), HA (A2095, Sigma), RENT1 (A300-038A, Bethyl), α-tubulin (T9026, Sigma), CNOT4 (12564-1-AP, Proteintech). Antibodies against GFP and RFP were a kind gift from Ramanujan Hegde. Secondary antibodies used were HRP-conjugated anti-Rabbit (170-6515, BioRad) and anti-Mouse (172-1011, BioRad), and HRP-conjugated Donkey anti-Goat (ab97110, Abcam).

siRNAs

Pre-designed Silencer Select siRNAs were ordered from ThermoFisher: control (scrambled 1), DCP1A (s31547), NEMF (137619), CNOT4 (s9631).

Mammalian cell culture

WT HEK293T cells were grown in Dulbecco's modified eagle medium (DMEM) with 10% FBS (Atlanta Biologicals, #S11550) and 2 mM L-glutamine (Invitrogen, #25030081). siRNA treatments were also performed according to manufacturer's instructions in a 6 well plate with 30 pmol of each siRNA, and analysed after 72 hours. Transient transfections were performed according to the manufacturer's protocol with 1 ug of reporter construct DNA, and fluorescence was measure after 24 hours.

Stable HEK293 cell lines were generated using Flp-In 293 T-Rex cells purchased from Thermo Fisher Scientific (USA) (RRID: CVCL_U427). Cell lines were grown in DMEM supplemented with 2 mM glutamine, 10% (w/v) FBS, 15 μg/ml Blasticidine S, and 100 μg/ml Zeocin. The open-reading frame to be integrated into the genomic FRT site was cloned into the pcDNA5/FRT/TO vector backbone and cell lines were generated according to the manufacturer's protocol. Briefly, the reporter construct was transfected together with pOG44 Flp-In recombinase in a 9:1 ratio using Trans-IT 293 transfection reagent (Mirus, USA) according to the manufacturer's instructions. 48 hours after transfection, 100 μg/ml Hygromycin B was used to select for cells that had undergone successful integration.

K562-dCas9-BFP-KRAB Tet-On cells (CMJ009) were grown in RPMI-1640 medium with L-Glutamine and HEPES (Lonza, #12-115F) supplemented with 10% Tet System Approved FBS, 100 units/mL penicillin and 100 μg/mL streptomycin (Invitrogen, #15140148). Cells were maintained at a confluency between 0.5-2 x$10^6$ cells/mL. All K562 cells harvested for FACS-based sorting were resuspended in RPMI 1640 without phenol red (Thermofisher Scientific, USA, 11835030) supplemented with 20% Tet System Approved FBS, 100 units/mL penicillin and 100 μg/mL streptomycin.

Lentivirus

Lentivirus was produced by co-transfecting HEK293T cells with two packaging plasmids (pCMV-VSV-G and delta8.9, Addgene #8454) and the corresponding transfer plasmid using TransIT-293 transfection reagent. 48 hours after transfection, the supernatant was collected, centrifuged and flash frozen. In all instances, virus was rapidly thawed prior to transfection. Virus for the genome-wide CRISPRi screen was generated using this method.

Virus generation for genome wide CRISPR knockout screen

HEK-293T cells were seeded at a density of 750,000 cells/mL in 20 mL viral production medium: IMDM (Thermo Fisher Scientific #1244053) supplemented with 20% inactivated fetal serum (GeminiBio #100-106). After 24 hours, media was changed to fresh viral production medium. At 32 hours post-seeding, cells were transfected with a mix containing 76.8 $\mu$L Xtremegene-9 transfection reagent (Sigma Aldrich #06365779001), 3.62 $\mu$g pCMV-VSV-G (Plasmid #8454, Addgene), 8.28 $\mu$g psPAX2 (Plasmid #12260, Addgene), and 20 $\mu$g sgRNA plasmid and Opti-MEM (Thermo Fisher Scientific #11058021) to a final volume of 1 mL. Media was changed 16 hours later to fresh viral production medium. At 48 hours after transfection, virus was collected and filtered through a 0.45 $\mu$m filter, aliquoted, and stored at -80 °C until use.

Generation of stable reporter cell lines

Reporter cell lines were generated by co-transfecting our control or NMD2 viral vectors along with a tet activator element into K562 wild type or K562 CRISPRi cell lines at one copy number per cell. Positive cells were isolated via FACS on a BD FACSAria2 and grown up to create monoclonal cell lines.

Flow cytometry analysis

For HEK293T experiments, RFP:GFP ratio was determined via FACS analysis of transfected or induced cells after 24 hours. Live HEK293T cells were first incubated with trypsin before collection, pelleted, and resuspended in 300 µL of PBS containing 1 µM Sytox Blue Dead Cell Stain (ThermoFisher, S34857) and analyzed on a Miltenyi Biotech MACSQuant VYB Flow Cytometer. K562 cells were infected at $0.5x10^6$ cells/mL. The media was supplemented with 8 µg/mL polybrene (Millipore Sigma, #107689-100G) and the lentivirus of interest was added to the well. The components were mixed by pipetting, and immediately spun down at 1000 g for 2 hours at 30C. After the spin, the culture was mixed. Expression of the gene of interest was tested after 48-72 hours. Data analysis for all flow cytometry experiments was performed using Python. In all instances, data is normalized to the control sample.

qPCR analysis

Relative mRNA levels were determined by quantitative PCR. Total cellular RNA was purified from cells using the RNeasy kit (74104, Qiagen), treated with DNase I (18068015, Invitrogen) and reverse transcribed using the SuperScript III First Strand Synthesis SuperMix (11752, Invitrogen), before being subjected to analysis on a StepOnePlus Real-Time PCR system. The relative expression ratios between sample cDNA levels were then analyzed, using primers against GFP and RFP, and the housekeeping gene HPRT1. Each set of primers was checked against a standard dilution curve, and the primer efficiencies were between 90 and 110%. The efficiencies were taken into account in the expression ratio calculation.

Inhibition of the ubiquitin-proteasome pathway

In Fig. 2, cells were treated with either 10 µM of the proteasome inhibitor MG132 (Calbiochem, 474790), or a DMSO control for 6 hours. To measure the RFP:GFP ratio at the protein level (sFig2), wild-type Flp-In 293 T-Rex cells were transiently transfected with FLAG-tagged versions of the reporter constructs. 18 hours later, reporter expression was induced with 1 µg/mL doxycycline and treated with either 10 µM of the E1 inhibitor MLN7243 or DMSO for 8 hours. Cells were harvested and lysed in 1% SDS. The lysates were normalized to GFP protein levels by serial dilutions and Western-blotting. The normalized lysates were analyzed by SDS-PAGE and Western-blotting using Anti-FLAG and Anti-GFP antibodies.

For Fig. 2B, our K562 CRISPRi NMD2 mononlocal cell line was induced with 1 µg/mL doxycycline for 6 hours and subsequently treated with 10 uM MG132 or DMSO. Cells were harvested and analyzed by flow cytometry on an Attune NxT Flow Cytometer.

To directly observe ubiquitination of RFP (Fig. 2C), cells were transiently transfected with FLAG-tagged versions of the reporter constructs in the presence and absence of HA-tagged ubiquitin and incubated for 42 hours. Cells were then treated with 10 uM MG132 for 6 hours and then harvested. Cells were resuspended in lysis buffer (50 mM Hepes pH 7.4, 100 mM KOAc, 2 mM $MgAc_2$, 1x Proteasome inhibitor, 1 mM DTT, 50 uM PR-619, 10 ug/mL Digitonin) and left on ice for 15 minutes. Mechanical lysis was performed with 10 strokes of a glass dounce and total samples taken. The amount of RFP in each sample was determined using a plate reader, and samples were normalized with HA-Ub-containing cell lysate to maintain the total protein concentration. SDS was then added to 1% final concentration, and the samples were boiled. They were then diluted with IP buffer (50 mM Hepes pH 7.4, 100 mM KOAc, 2 mM $MgAc_2$, 1% Triton) to a final concentration of 0.1% SDS. Samples were immunoprecipitated with Anti-FLAG M2 affinity resin (Millipore-Sigma) and eluted with SDS. Analysis was performed by Western blot.

#### CRISPRi knockdown screen

The genome-scale CRISPRi screen was performed similarly to previously described screens (Gilbert et al., 2014, Horlbeck et al., 2016). The hCRISPRi-v2 compact library (containing 5 sgRNAs per gene, Addgene pooled libray #83969) was transduced in duplicate into 330 million K562-CMJ009-NMD-2 cells at MOI < 1 (percentage of transduced cells 48 hours after infection as measured by BFP positive cells: 20%-40%). Cells were grown in 1L of media in 1L spinner flasks (Bellco, SKU: 1965-61010) for the duration of the screen. 48 hours after spinfection, cells were selected with 1 mg/mL puromycin for 3 days, after which transduced cells constituted 80-95% of the population. After a 36 hours recovery, cells were induced with 1 µg/mL doxycycline for 24 hours and sorted on a FACS AriaII Fusion Cell Sorter. The cells were maintained at $0.5x10^6$ cells/mL for the duration of the screen. Therefore, cells were diluted daily to $0.5x10^6$ cells/mL except during and after puromycin treatment, when dilutions depended on the health of the culture and were more conservative due to puromycin-caused cell. This ensured that the culture was maintained at an average coverage of more than 1000 cells per sgRNA for the whole screen.

Cells with high BFP (transduced cells) and with both GFP and RFP signal (successfully induced) were gated. Cells were sorted according to the RFP:GFP ratio of this population.
Around 40 million cells with either the highest (30%) and the lowest (30%) RFP:GFP ratio were collected, pelleted and flash-frozen. Genomic DNA was purified using the Nucleospin Blood XL kit (Takara Bio, #740950.10) and amplified with barcoded primers by index PCR. The library (~264 bp) was purified using SPRIbeads (Bulldog Bio, CNGS005), its concentration measured by Qubit fluorometer (Invitrogen) and its integrity checked by Agilent 2100 Bioanalyzer. Samples were analyzed using an Illumina HiSeq2500 high throughput sequencer. Sequencing reads were aligned to the CRISPRi v2 library sequences, counted and quantified (Horlbeck et al., 2016). Generation of negative control genes and calculation of phenotype scores and Mann-Whitney p-values was performed as described previously (Gilbert et al., 2014; Horlbeck et al., 2016). Gene-level phenotypes and counts are available in Supplementary Table 1.

#### K562 genome-wide CRISPR knockout screen

A genome-wide lentiviral sgRNA library in a Cas9-containing vector (unpublished, Supplementary Table 3) was used to transduce 500M K562 cells. All other conditions were identical to those used for the CRISPRi KD screen. Cells were induced either at 7, 9, or 11 days with 1 µg/mL doxycycline for 24 hours and sorted on a FACS Aria II Fusion cell Sorter. Data was processed using the pipeline described above and validated by analysis using MAGeCK (Li et al., 2014). Gene-level phenotypes and counts are available in Supplementary Table 2.

For extraction of genomic DNA, QIAamp DNA Blood Maxiprep Kit (Qiagen) was used according to manufacturer's instructions with the following modifications: 500 $\mu$L of a 10 mg/mL solution of ProteinaseK (Fisher #) in water was used in place of QIAGEN Protease; incubation with ProteinaseK and Buffer AL was performed overnight; centrifugation steps after Buffer AW1 and AW2 were performed for 2 min and 5 min, respectively; gDNA was eluted for 5 min using 1 mL of water preheated to 70 °C, followed by centrifugation for 5 min. gDNA concentration was determined using the Qubit dsDNA HS Assay kit (Thermo Fisher Scientific #Q32851).

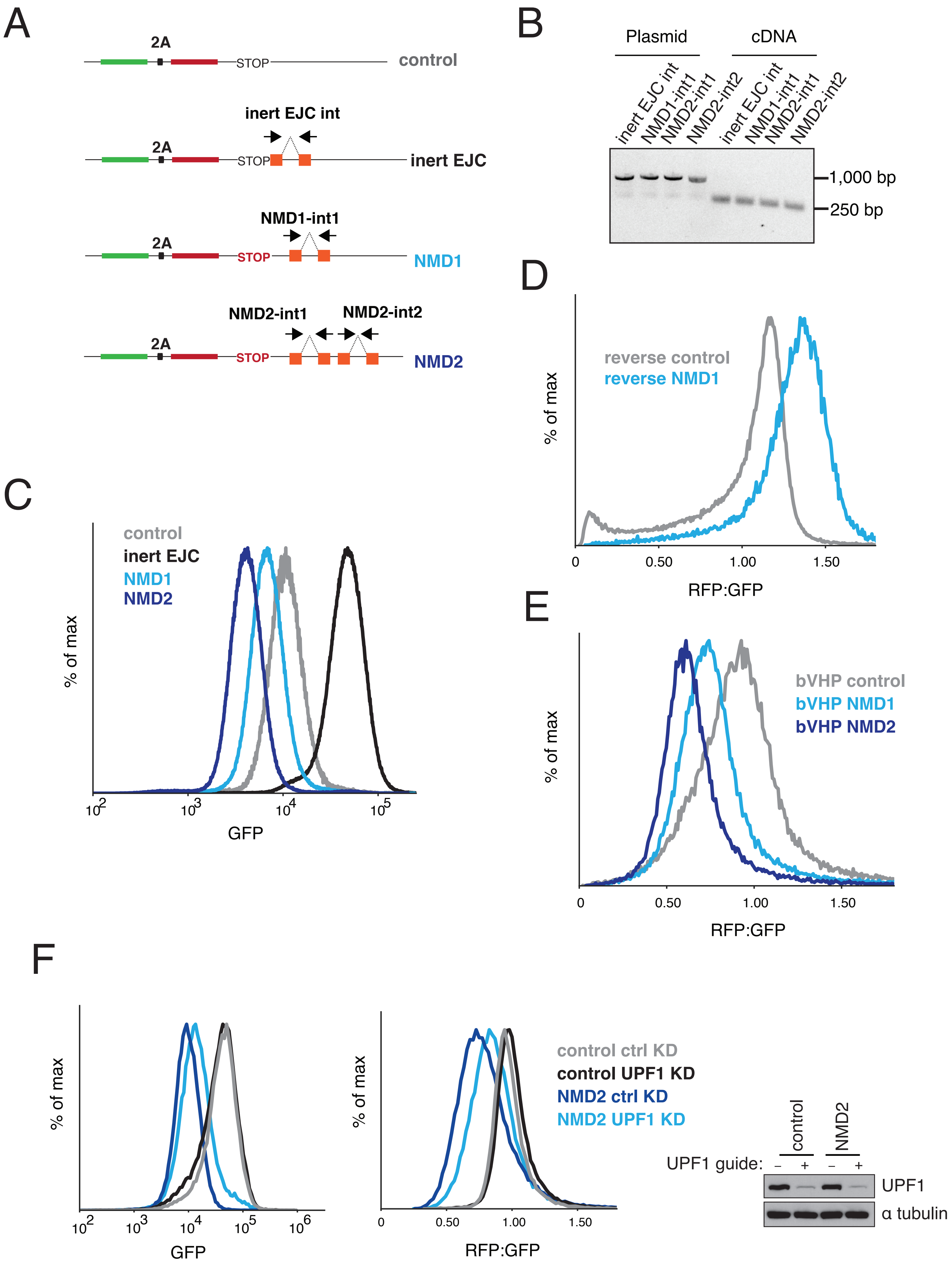
A
2A
STOP
control
inert EJC int
inert EJC
NMD1-int1
NMD1
NMD2-int1
NMD2-int2
NMD2
B
Plasmid
cDNA
inert EJC int
NMD1-int1
NMD2-int1
NMD2-int2
1,000 bp
250 bp
C
control
inert EJC
NMD1
NMD2
% of max
GFP
D
reverse control
reverse NMD1
% of max
RFP:GFP
E
bVHP control
bVHP NMD1
bVHP NMD2
% of max
RFP:GFP
F
% of max
GFP
RFP:GFP
control ctrl KD
control UPF1 KD
NMD2 ctrl KD
NMD2 UPF1 KD
control
NMD2
UPF1 guide:
UPF1
α tubulin

sFig.1

**sFig 1. Characterization of the NMD reporter constructs.**

(A) Reporter design as depicted and described in Fig 1A. Orange boxes represent the sequence from the β-globin gene, with the dashed lines representing constitutive splice sites. For NMD1 and NMD2, the first intron was inserted ~200 bp from the stop codon (int1). The NMD2 reporter contained a second β-globin splice site (int2) that resulted in further suppression of the mRNA levels. To account for changes in translation associated with the deposition of an exon junction complex (EJC) following splicing, we created an additional control with the β-globin intron only 12 bp from the stop codon, an insufficient distance for recognition by NMD machinery. Black arrows indicate the position of primers used to check efficient splicing in (B). (B) Both β-globin introns were efficiently spliced in their ectopic context. Either plasmid DNA or the corresponding cDNA for the indicated reporters was amplified by PCR using primer pairs that span the introns. Upon splicing, the size of the expected fragment decreases from 1 kb to ~250 bp. (C) GFP channel of samples analyzed by flow cytometry in Figure 1A. (D) Schematic of reporter in which RFP and GFP order is reversed and the corresponding analysis of these reporters by flow cytometry. The enhanced degradation of the 3' moiety is independent of fluorescent protein identity. (E) A hydrophilic linker was inserted between the RFP and stop codon in the indicated reporters to ensure the RFP was fully emerged from the ribosome at the stop codon. The RFP:GFP ratio of these reporters is displayed as a histogram. (F) K562 cells containing CRISPRi machinery stably expressing the NMD2 reporter or the control reporter under an inducible promoter were infected with dual guides against UPF1 or a non-targeting control. After 8 days, cells were induced with doxycycline, and harvested after 24 hours for analysis by flow cytometry and Western blotting.

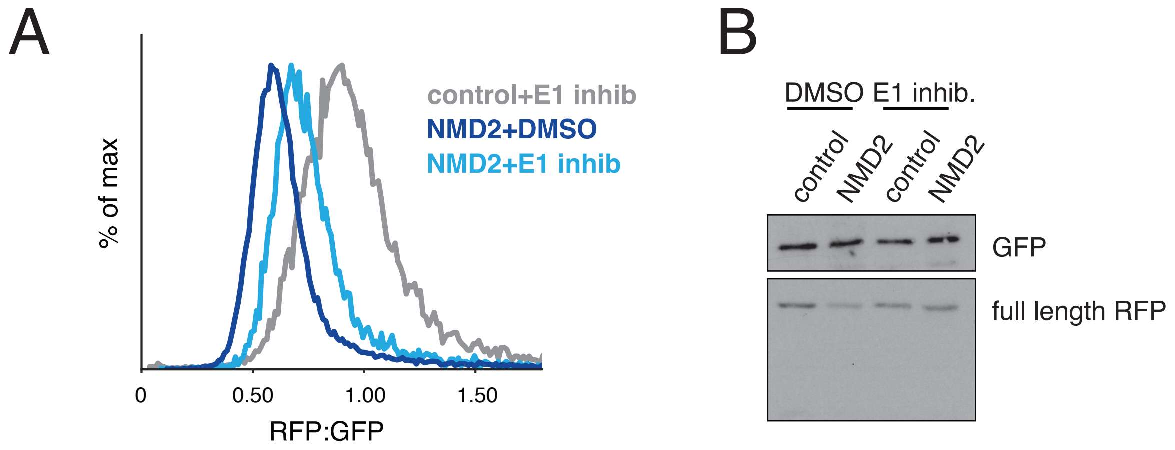


**sFig 2. NMD-linked nascent protein degradation is dependent on the ubiquitin-proteasome system.** (A) Wild-type HEK293T cells were transiently transfected with either control or NMD2 reporters. After 16 hours, the cells were treated for 8 hours with either 10 μM MLN7243 or a matched DMSO control. Cells were then harvested and analyzed by flow cytometry. (B) Cells were treated as in (A), but were lysed in 1% SDS after MLN7243 treatment. The lysates were boiled and subjected to SDS-PAGE and Western blotting. Samples were normalized to GFP to control for RNA degradation. No RFP degradation products were observed.

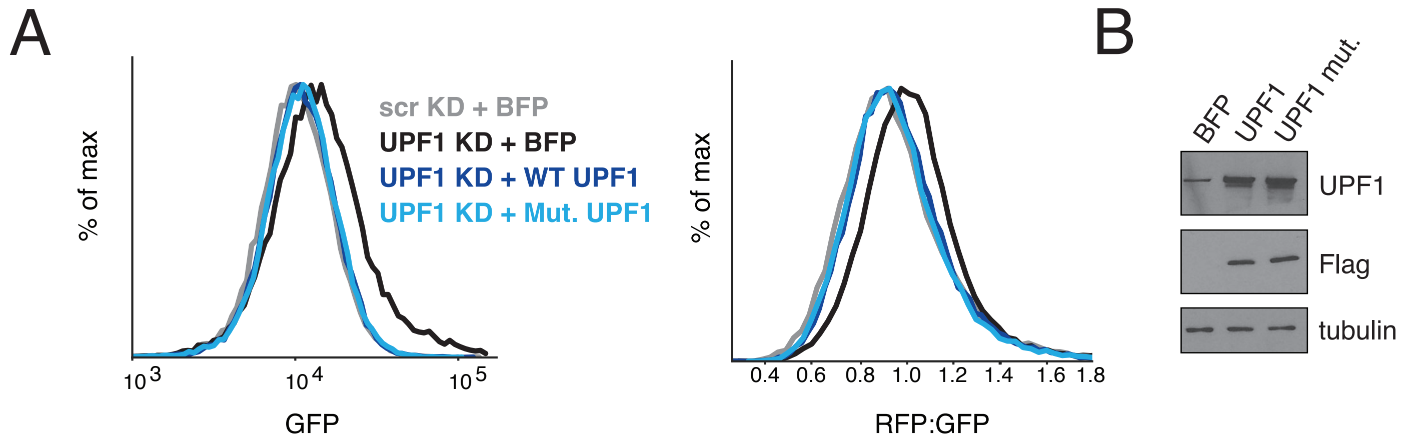


**sFig 3. The role of E3 ligases in NMD nascent chain degradation.**
K562 cell lines stably expressing CRISPRi machinery and the inducible NMD2 reporter were constructed to stably express one copy of either BFP, a FLAG-conjugated wild-type UPF1, or a FLAG-conjugated mutant UPF1 (S134A, N148A, T149A) with disruptions that abolish association with E2 conjugating enzymes. WT or mutant UPF1 was separated from BFP by a viral P2A sequence, allowing us to use BFP as a proxy for UPF1 infection. These cells were then infected with dual guides targeting UPF1 or a non-targeting control. After 8 days of knockdown, the NMD2 reporter was induced with doxycycline for 24 hours, after which cells were harvested and analyzed by flow cytometry (A) and Western blotting (B).

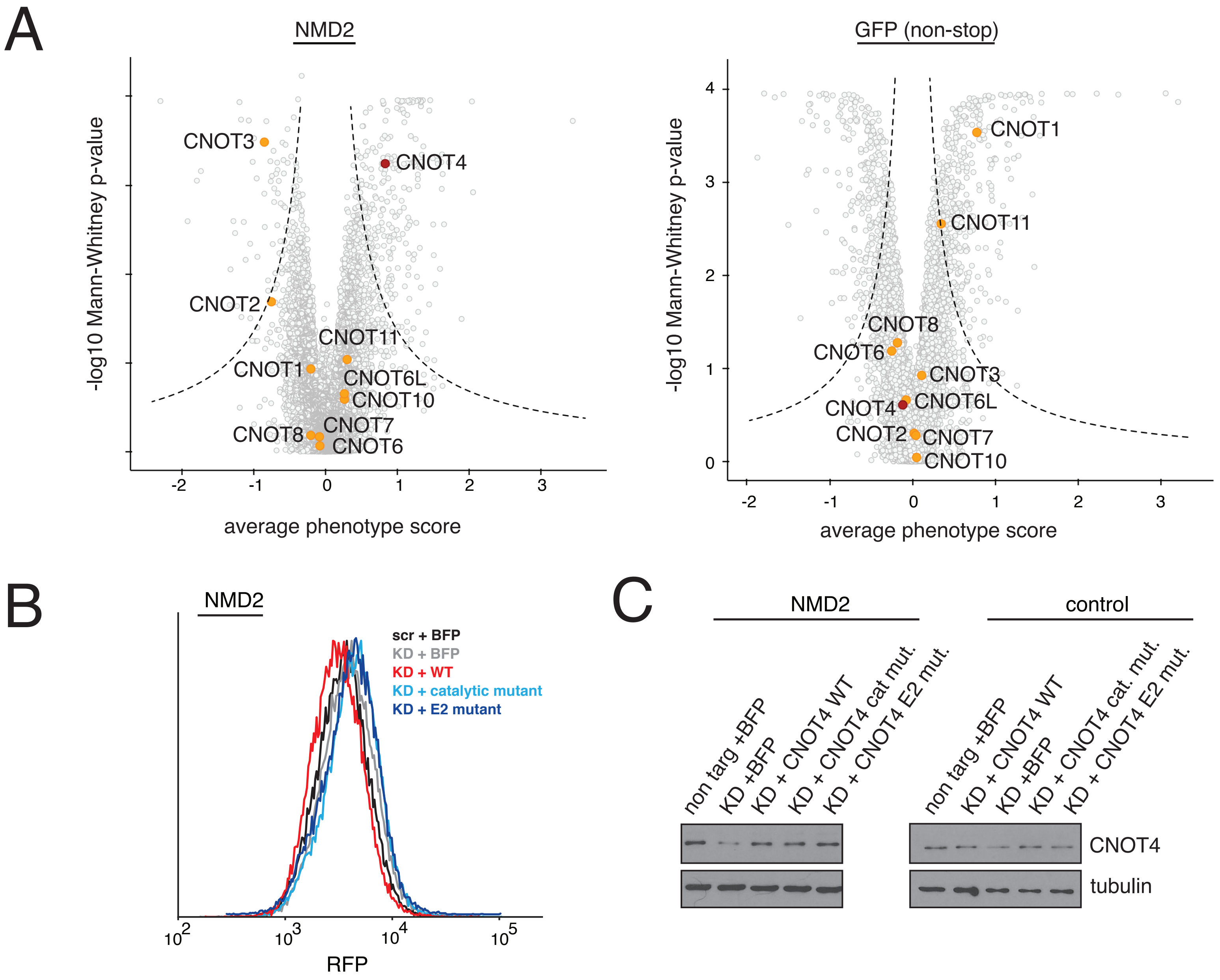


**sFig 4. Factors involved in NMD-coupled protein quality control.**
(A) Volcano plots of CRISPRi screens with our NMD2 reporter or GFP non-stop reporter (Hickey et al., 2020). In orange are highlighted components of the CCR4-NOT complex, except for CNOT4 which is shown in red. CNOT4 has a significant effect in the NMD2 reporter screen, increasing the level of RFP relative to GFP. (B) RFP channel of samples analyzed by flow cytometry in Figure 5C. (C) Expression levels of wild type and mutant CNOT4 in the experiment displayed in Fig. 6B as determined by Western blotting.
